# The Human Chk1 Inhibitor CHIR-124 Shows Multistage Activity Against the Human Malaria Parasite *Plasmodium falciparum* via Polypharmacological Inhibition of *Pf*Ark1 and Hemozoin Formation

**DOI:** 10.1101/2025.07.17.664511

**Authors:** Kathryn J. Wicht, John G. Woodland, Larnelle F. Garnie, Henrico Langeveld, Dale Taylor, Luiz C. Godoy, Charisse Flerida A. Pasaje, Mariana Laureano de Souza, Jair L. Siqueira-Neto, Sonja Ghidelli-Disse, Maria Jose Lafuente-Monasterio, Francisco-Javier Gamo, Dina Coertzen, Janette Reader, Mariëtte van der Watt, Jessica L. Bridgford, Gareth Girling, Rachael Coyle, Christian Scheurer, Sergio Wittlin, Arne Alder, Tim-Wolf Gilberger, Marcus C.S. Lee, Till S. Voss, Elizabeth A. Winzeler, David A. Fidock, Jacquin C. Niles, Lyn-Marié Birkholtz, Lauren B. Coulson, Kelly Chibale

## Abstract

The high burden of malaria and growing resistance to frontline antimalarials demand new drug target combinations with reduced propensities for conferring parasite resistance. An attractive approach for circumventing antimalarial drug resistance is target repurposing in which known drugs that act through protein targets of human origin that are also active against the human malaria parasite *Plasmodium falciparum* are exploited to identify novel antimalarial drug targets. Here we show that the human checkpoint kinase 1 (Chk1) inhibitor CHIR-124 is active *in vitro* against both drug-sensitive and drug-resistant asexual blood stage parasites and competitively binds to several *Plasmodium* kinases. The compound also shows moderate activity against both the liver and gametocyte forms of the parasite. Further target investigation of CHIR-124 via conditional knockdown experiments confirmed that *P. falciparum* Aurora-related kinase 1 (*Pf*Ark1) is implicated in its parasiticidal activity. Notably, CHIR-124 also inhibits β-hematin (synthetic hemozoin) formation and causes a dose-dependent increase in free heme that correlates with inhibition of parasite growth. These findings suggest that polypharmacology is involved in the activity of CHIR-124 against *P. falciparum* via the dual inhibition of *Plasmodium Pf*Ark1 and hemozoin formation, both essential for parasite proliferation. This is further supported by *in vitro* drug combination experiments, morphological studies and resistance generation attempts. This study validates the feasibility of dual *Plasmodium* kinase/hemozoin formation inhibitors active against resistant strains with decreased resistance risks in the fight against malaria.

## Introduction

The mosquito-borne disease of malaria is caused by protozoan parasites of the *Plasmodium* genus, of which *P. falciparum* is the most pervasive and deadly to humans.^1^ *P. falciparum* parasites have a complex life cycle involving over ten different morphological forms, with the asexual blood stage (ABS) in the human host giving rise to clinical pathology in malaria patients.^2, 3^ Hypnozoites (liver stage) and gametocytes (transmission stage) are also biologically relevant life cycle stages within the human host. However, rapidly reducing the parasite burden at the ABS stage is the primary objective of most therapeutic antimalarials.^4^ Drugs active against the ABS are categorized by the Medicines for Malaria Venture (MMV) Target Candidate Profiles (TCPs) as TCP1.^5^ Nevertheless, for malaria eradication, strategies that break the cycle of disease transmission need to be considered.^6,7^ TCP1 molecules should ideally be paired with a partner drug also targeting the liver stage hypnozoites for prophylactic treatment (TCP4) or the transmission-blocking stage gametocytes (TCP5) in addition to the ABS.^6, 8^

Unfortunately, malaria mortality and incidence rates have increased since 2015, largely due to the slowed progress of the World Health Organization (WHO) Global Malaria Programme and further setbacks from the Covid-19 pandemic.^9^ The WHO reported an estimated 282 million malaria cases and 610,000 malaria deaths worldwide in 2024, and highlighted that 94% of the disease burden is in sub-Saharan Africa.^10^ With partial drug resistance to essential frontline regimens such as the artemisinin (ART) derivatives and their partner drugs on the rise, this is a critical time for discovering new antimalarials.^11^ Historically, developing such therapies has been an immensely time-consuming and expensive task, with the discovery phase requiring that drug candidates meet strict criteria to progress to clinical development.^12, 13^ This has led to high attrition rates and prompted the desire for more efficient strategies for identifying new drug targets and drug molecules, particularly those with a low risk for generating resistance in parasite populations.^14^

One approach to circumvent antimalarial drug resistance is the identification of novel *Plasmodium* drug targets on which to base target-driven antimalarial drug discovery. To this end, target repurposing takes advantage of previously approved compounds that have already been through clinical trials and safety studies that act through protein targets of human origin by evaluating their efficacy for new indications.^15, 16^

Within the context of malaria, identifying target-focused molecules for target repurposing is facilitated in cases where there is an ortholog target protein family in *Plasmodium*. For this purpose, human kinase inhibitors active against parasites provide a promising avenue for further investigation as putative *Plasmodium* kinase inhibitors.^17^ Several *Plasmodium* kinases have been identified as favorable antimalarial targets, given their druggability and essentiality during more than one stage of the parasite life cycle.^18^ Kinases are central to signal transduction pathways and play critical physiological roles in cell growth and proliferation.^19^ Given the pivotal role of kinase inhibitors in non-communicable diseases, a wealth of structural data supports the design and optimization of kinase inhibitors.^20^

Only one *Plasmodium* kinase inhibitor, MMV390048, which targets phosphatidylinositol 4-kinase type III beta (*Pf*PI4K), has reached the clinical stage of development for malaria so far.^21^ Nevertheless, multiple opportunities exist for exploitation of this class of drug targets. Given the conserved nature of the ATP-binding site across the kinase superfamily, kinase inhibitors have the potential to display polypharmacology, which is advantageous for minimizing the generation of resistance.^18, 22^ Although selectivity for *Plasmodium* relative to human kinases is a key challenge for kinase-focused malaria drug discovery, identifying compounds that bind to *Plasmodium* kinases is an appealing approach, not only for finding new scaffolds with antimalarial activity but also for discovering kinase-inhibiting tool compounds.

Here we disclose the identification of the human checkpoint 1 (Chk1) inhibitor CHIR-124 from a phenotypic ABS screen of clinical or pre-clinical anticancer human kinase inhibitors. We show that CHIR-124 possesses multistage activity against *P. falciparum* as well as retained potency against drug-resistant strains. Target identification via *Plasmodium* kinases captured on Kinobeads as well as biochemical, conditional knockdown, intracellular heme fractionation, morphological and drug combination studies reveal that polypharmacology is involved in the mode of action of CHIR-124.

## Results

### CHIR-124 Identified via Kinobead Assays with Multistage Antiplasmodial Activity

A set of clinical human kinase inhibitors in clinical or pre-clinical development, selected based on their target diversity for the treatment of cancer, was screened against *P. falciparum*. The human Chk1 inhibitor, CHIR-124 (GSK3210608, **Figure 1A**), was identified with activity <1 µM against *Pf*3D7 (wild-type, drug sensitive) parasites using the previously reported [^3^H]-hypoxanthine incorporation assay.^23^ In humans, Chk1 is a serine/threonine kinase that regulates the G^2^/M cell cycle checkpoints and delays cell cycle progression in response to DNA damage.^24^ Disrupting Chk1-mediated checkpoints renders tumor cells more susceptible to DNA damage and is therefore a convenient target of chemotherapy for cancers.^25^ CHIR-124 has antitumor properties when used in combination with topoisomerase I poisons.^26^

**Figure 1.**
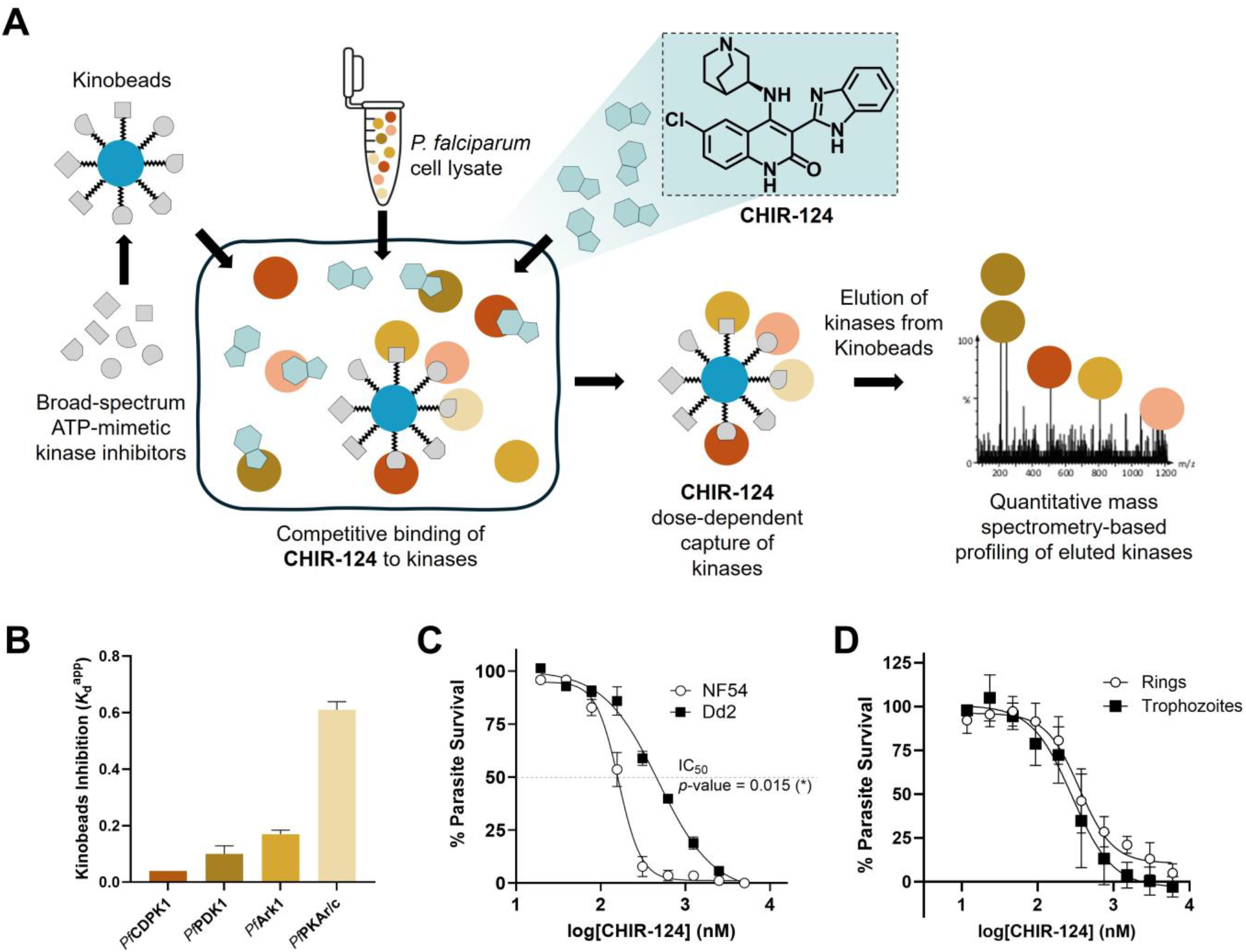
(**A**) The human kinase inhibitor CHIR-124 competitively binds to *Plasmodium* kinases captured on Kinobeads by broad spectrum kinase inhibitors immobilized on solid support beads. Eluted kinases are quantified in a dose-dependent manner via mass spectrometry. (**B**) Competitive binding for the top *Plasmodium* kinases with lowest apparent dissociation constant (*K*_*d*_^app^) in micromolar competed off Kinobeads by CHIR-124. Data and error bars represent the mean and SD from two independent experiments. (**C**) Dose-response profiles of CHIR-124 against the wild-type *Pf*NF54 strain vs the multidrug-resistant *Pf*Dd2 strain. Data are plotted as the mean with error bars representing the SEM where N,n=3,2. IC_50_ values are significantly different with \**p*<0.05 via an unpaired two-tailed *t*-test. (**D**) Stage specificity of CHIR-124 in *Pf*NF54 showing dose response in rings and trophozoites. Data from rings vs trophozoites represent the mean ± SD from N,n=2,3-4 and are not significantly different.

To identify potential kinase targets of CHIR-124 in *Plasmodium*, the compound was screened against 75 *Plasmodium* kinases captured onto Kinobeads from ABS parasite lysates using a competitive pulldown assay (**Figure 1A**).^27, 28^ A total of eight *Plasmodium* kinases were competed off the Kinobeads by CHIR-124 at 10 µM with a log_2_ fold-change below the cutoff of -1 relative to the vehicle control (**Supplementary Figure S1** and **Supplementary Table S1 Excel**). Of these, five showed half-maximal inhibitory concentration (IC_50_) values below 1 µM with apparent dissociation constants *K*_d_^app^ below 0.7 µM. They included the calcium-dependent protein kinase 1 (*Pf*CDPK1, PF3D7_0217500, *K*_d_^app^ 0.04 µM), 3-phosphoinositide-dependent protein kinase 1 (*Pf*PDK1, PF3D7_1121900, *K*_d_^app^ 0.1 µM), Aurora-related kinase 1 (*Pf*Ark1, PF3D7_0605300, *K*_d_^app^ 0.17 µM) and the cAMP-dependent protein kinase regulatory/catalytic subunits (*Pf*PKAr/c, PF3D7_1223100/PF3D7_0934800, *K*_d_^app^ 0.61 µM) (**Figure 1B, Supplementary Table S1**). Given that *Plasmodium* kinases are generally expressed across multiple stages of the parasite life cycle, albeit to differing degrees,^29^ the potential of CHIR-124 to inhibit several *Plasmodium* kinases prompted us to evaluate its activity across the three different host life cycle stages.

For the dose-dependent growth inhibition of *P. falciparum* by CHIR-124 across the life cycle stages, we profiled CHIR-124 against *in vitro* cultures of the ABS, both immature and late-stage gametocytes, and the liver stage. These assays confirmed the sub-micromolar activity of CHIR-124 against ABS parasites in the drug-sensitive strain (**Figure 1C, Table 1** and **Supplementary Table S2**). Moderate activity was also observed against immature and late-stage gametocytes, in the low micromolar range, as well as against liver stages, in the sub-micromolar range (**Table 1** and **Supplementary Table S3**). Furthermore, CHIR-124 displayed roughly 14-fold selectivity for liver stage parasites relative to the HepG2 mammalian liver cell line (**Table 1** and **Supplementary Table S3**). The multistage activity of CHIR-124 suggests that at least one of its intracellular targets is biologically crucial at each life cycle stage tested.

**Table 1.**
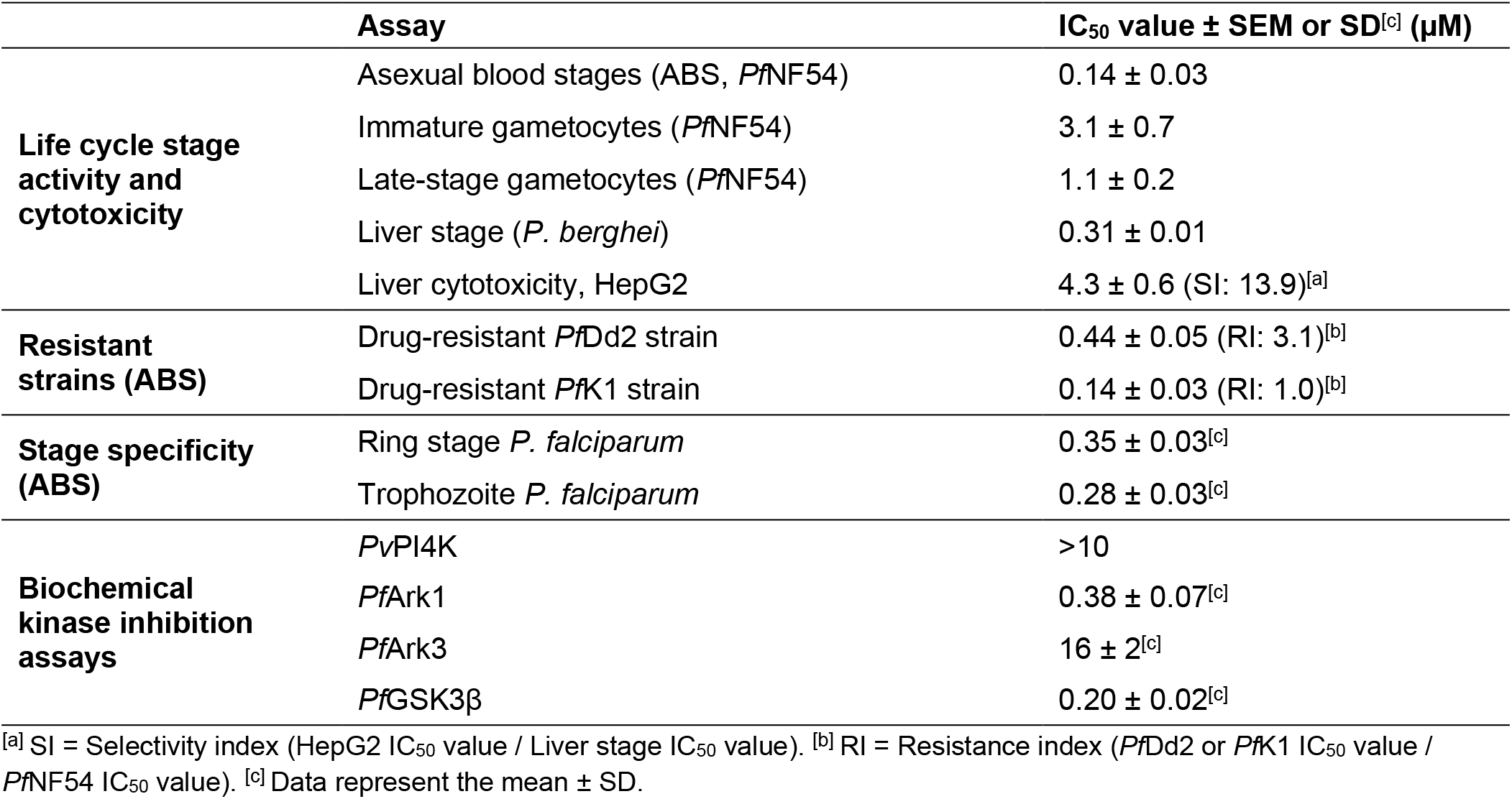
Life cycle activity, cross-resistance, stage specificity and biochemical inhibition data for CHIR-124.

|  | Assay | IC<sub>50</sub> value ± SEM or SD<sup>[c]</sup> (μM) |
| --- | --- | --- |
| Life cycle stage activity and cytotoxicity | Asexual blood stages (ABS, *Pf*NF54) | 0.14 ± 0.03 |
|  | Immature gametocytes (*Pf*NF54) | 3.1 ± 0.7 |
|  | Late-stage gametocytes (*Pf*NF54) | 1.1 ± 0.2 |
|  | Liver stage (*P. berghei*) | 0.31 ± 0.01 |
|  | Liver cytotoxicity, HepG2 | 4.3 ± 0.6 (SI: 13.9)<sup>[a]</sup> |
| Resistant strains (ABS) | Drug-resistant *Pf*Dd2 strain | 0.44 ± 0.05 (RI: 3.1)<sup>[b]</sup> |
|  | Drug-resistant *Pf*K1 strain | 0.14 ± 0.03 (RI: 1.0)<sup>[b]</sup> |
| Stage specificity (ABS) | Ring stage *P. falciparum* | 0.35 ± 0.03<sup>[c]</sup> |
|  | Trophozoite *P. falciparum* | 0.28 ± 0.03<sup>[c]</sup> |
| Biochemical kinase inhibition assays | *Pv*PI4K | >10 |
|  | *Pf*Ark1 | 0.38 ± 0.07<sup>[c]</sup> |
|  | *Pf*Ark3 | 16 ± 2<sup>[c]</sup> |
|  | *Pf*GSK3β | 0.20 ± 0.02<sup>[c]</sup> |

<sup>[a]</sup> SI = Selectivity index (HepG2 IC<sub>50</sub> value / Liver stage IC<sub>50</sub> value). <sup>[b]</sup> RI = Resistance index (*Pf*Dd2 or *Pf*K1 IC<sub>50</sub> value /
*Pf*NF54 IC<sub>50</sub> value). <sup>[c]</sup> Data represent the mean ± SD.

We also profiled CHIR-124 against ABS drug-resistant strains, including the Southeast Asian CQ-resistant lines *Pf*Dd2 and *Pf*K1, as well as the ART-resistant line *Pf*Cam3.II and the ART- and piperaquine-resistant *Pf*RF7 line. The growth of all strains was inhibited by sub-micromolar concentrations of CHIR-124 but, interestingly, an increased IC_50_ value (∼3-fold less active) was observed against *Pf*Dd2 relative to the *Pf*NF54 wild-type strain (*p*-value=0.015, **Figure 1C**), whereas against the *Pf*K1 strain the IC_50_ value was indistinguishable (**Table 1** and **Supplementary Table S2**). This 3-fold increase in IC_50_ against the *Pf*Dd2 strain could indicate that the compound is marginally susceptible to multidrug resistance-mediated mechanisms (e.g. amplification of *pfmdr1*) as these are absent in the *Pf*K1 strain.^30^ Furthermore, CHIR-124 showed differences in dose-response curve profiles against the ART-resistant strains *Pf*Cam3.II and *Pf*RF7 relative to *Pf*NF54, where shallow or biphasic curves were observed (**Supplementary Figure S2** and **Supplementary Table S2**). These findings suggest that, while CHIR-124 remains active against drug-resistant strains, it is marginally cross-resistant with CQ and piperaquine. To further probe for cross-resistance, a pool of 45 DNA-barcoded resistant mutant lines (AReBar assay)^31, 32^ was exposed to 3×IC_50_ concentrations of CHIR-124 (**Supplementary Table S4)**. No mutant strain outgrowth was observed after 14 days, suggesting negligible cross-resistance against known mutants (**Supplementary Figure S3**). We further investigated the ABS activity of CHIR-124 in rings versus trophozoites (**Figure 1D**) and observed minimal differences, suggesting that CHIR-124 targets processes of importance to both these stages.

### Conditional Knockdown and Biochemical Studies Reveal PfArk1 as a Target of CHIR-124

We then further investigated each of the *Plasmodium* kinase targets competitively inhibited by CHIR-124 via the Kinobead studies in pursuit of the bona fide intracellular targets. The kinase with the lowest *K*_d_^app^, *Pf*CDPK1 (**Figure 1B**), is unlikely to be responsible for antiplasmodial activity based on previous studies showing that *Pf*CDPK1 is non-essential for ABS parasite survival.^33, 34^ Hence, we employed conditional knockdown (cKD) susceptibility assays with knockdowns of the remaining candidate kinases: *Pf*PDK1, *Pf*PKAc and *Pf*Ark1. We also included the cKDs of *Pf*Ark2 and the negative control *Pf*PI4K for comparison.

Two knockdown approaches were used. The first, for *Pf*PDK1, was based on tagging the endogenous *pfpdk1* gene with *gfp* fused to the *dd* sequence to generate the *Pf*PDK1-GFPDD protein in *Pf*NF54 WT parasites.^35^ CHIR-124 was profiled against cultures under *Pf*PDK1-GFPDD-depleting conditions (–Shield-1) versus *Pf*PDK1-GFPDD-stabilizing conditions (+Shield-1). Similar CHIR-124 IC_50_ values were obtained regardless of *Pf*PDK1 protein expression, indicating negligible interaction between CHIR-124 and *Pf*PDK1 in whole cells (**Supplementary Table S5**).

The second cKD approach, used for *Pf*Ark1, *Pf*Ark2, *Pf*PKAc and *Pf*PI4K, relies on tetracycline transcriptional regulator (TetR)-binding RNA aptamers, whereby binding to the aptamer is inhibited by increasing the concentration of anhydrotetracycline (aTc) to allow downstream translation of mRNA and protein synthesis.^36^ Normal levels of kinase expression are achieved when ≥50 nM aTc is added to the culture prior to assay setup. The dose-response activity of the cell line fully expressing the target (50 nM or 500 nM aTc) is then compared to that of the culture with low (1 nM) aTc in which expression of the kinase is repressed. Data showing the effect of the kinase knockdown on untreated parasite viability is shown for *Pf*Ark1, *Pf*Ark2 and *Pf*PKAc in **Supplementary Figure S4**. Studies with CHIR-124 treatment revealed that the *Pf*Ark1 cKD line where expression of *Pf*Ark1 is reduced using 1 nM aTc is significantly more sensitive to CHIR-124 by 25-fold (*p*-value=0.011) relative to the line using 500 nM aTc that fully expresses the kinase (left-shift of the dose-response curve, **Figure 2A**). We noted that this 25-fold shift is larger than that of the known *Pf*Ark1 inhibitor, hesperadin, which showed a 7-fold shift in the *Pf*Ark1 cKD experiment.^37^ Only minor shifts were observed with cKD lines *Pf*PKAc and *Pf*Ark2 of 3.6-fold and 2.9-fold, respectively, similar to those observed with the *Pf*PI4K negative control line (**Supplementary Figure S4A-D** and **Supplementary Table S5**), suggesting that these kinases do not play significant roles in the antiplasmodial activity of CHIR-124.

**Figure 2.**
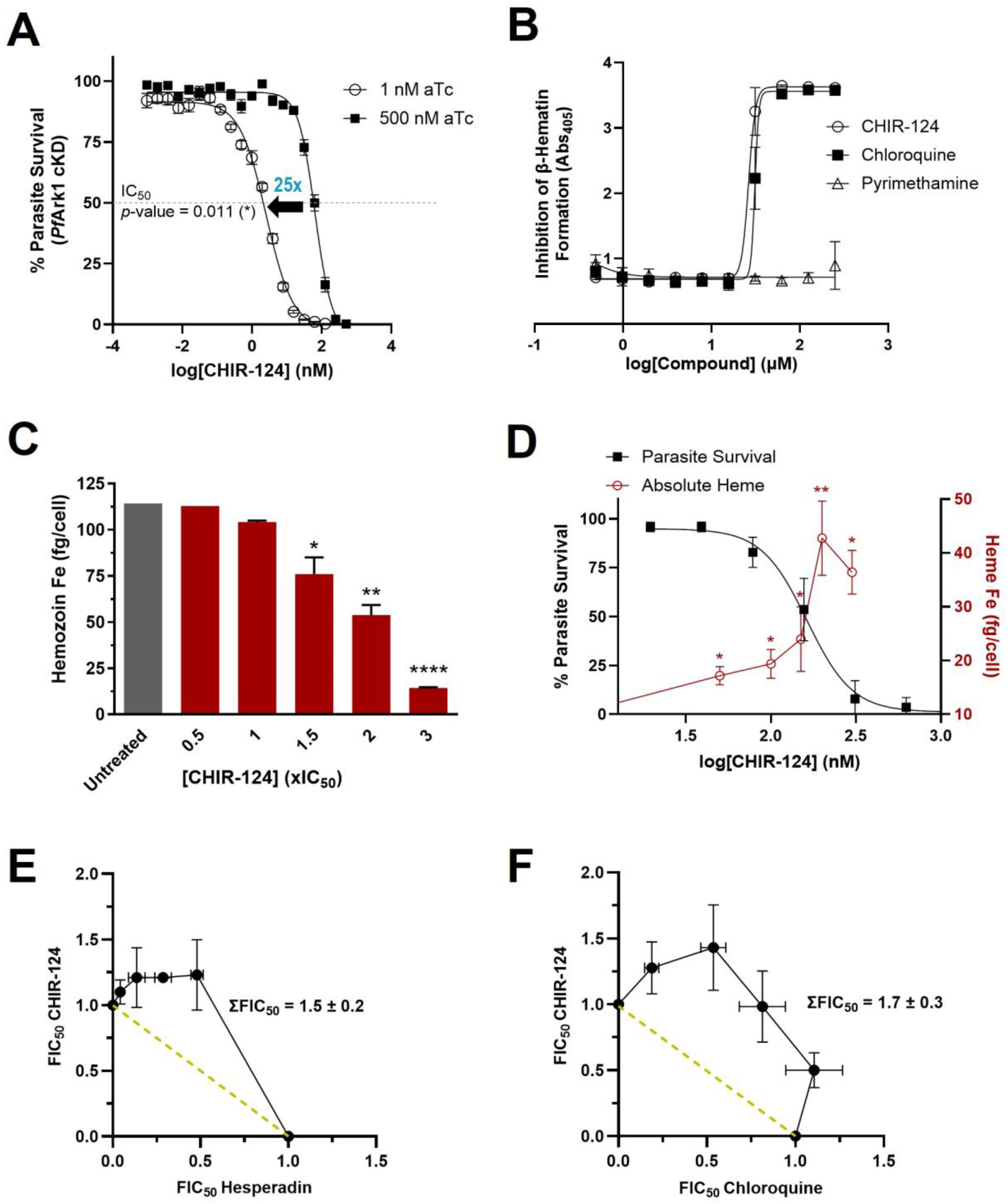
Genetic and phenotypic target exploration studies for CHIR-124. (**A**) The *Pf*Ark1 cKD line exposed to CHIR-124 under low (1 nM aTc) or high (500 nM aTc) conditions of protein expression. Data represents mean ± SEM where N,n=3,3. IC_50_ values are significantly different with \**p*<0.05 via an unpaired two-tailed *t*-test. (**B**) β-hematin inhibition activity of CHIR-124 in the extracellular assay detergent-based biomimetic assay. Data represent mean ± SD (n=3). (**C**) Dose-dependent changes in the heme Fe levels from intracellularly-extracted fractions of hemozoin under CHIR-124 treatment (representative from N,n=3,3). Error bars represent ± SD for technical triplicates with an unpaired two-tailed *t*-test, where *p*-values are shown for \**p*<0.05; \*\**p*<0.01; \*\*\**p*<0.001; \*\*\*\**p*<0.0001. (**D**) Parasite survival overlaid with the intracellularly-extracted fractions of free heme under CHIR-124 treatment. Significance determined as per panel (**C**). The correlation observed between the increasing heme levels with decreasing parasite survival indicates that hemozoin inhibition is a contributing mechanism of action of CHIR-124. Combination studies via fixed-ratio isobologram analysis showing an antagonistic effect on *Pf*NF54 ABS parasites for CHIR-124 with (**E**) the *Pf*Ark1 inhibitor hesperadin or (**F**) the hemozoin formation inhibitor chloroquine. Data represent mean fractional 50% inhibitory concentration (FIC_50_) values ± SEM from three biological replicates each with 2–3 technical replicates. The mean sum of the FIC_50_ values (ΣFIC_50_) ± SEM is also shown for each combination where ΣFIC_50_ >1.2 indicates antagonism.

CHIR-124 was also screened against available recombinant *Plasmodium* kinases (Luceome Biotechnologies) via the competition binding KinaseSeeker™ assay.^38^ In this assay format, each kinase is expressed in cell-free lysate without further purification and displacement of a probe from the ATP binding site is measured. Using this approach, further validation of the interaction between CHIR-124 and *Pf*Ark1 was provided (*Pf*Ark1 IC_50_ 0.38 µM, **Table 1**), albeit at higher concentrations than the *Pf*Ark1 inhibitor hesperadin (*Pf*Ark1 IC_50_ 0.002 µM).^37^ In contrast, negligible inhibition of *Pf*Ark3 was observed (IC_50_ 16 µM), corroborating the fact that this kinase was not identified as a putative target in the Kinobead experiment. However, the most potent biochemical activity for CHIR-124 against this kinase panel was seen for glycogen synthase kinase 3 beta (*Pf*GSK3β, PF3D7_0312400) with an IC_50_ of 0.2 µM (**Table 1** and **Supplementary Figure S5**). Notably, *Pf*GSK3β was also identified in the Kinobead experiments above, albeit with an IC_50_ of 3 µM (**Supplementary Figure S1**). The discrepancy in CHIR-124 IC_50_ values between the KinaseSeeker™ and Kinobead assays is perhaps not unexpected, as they operate in fundamentally different systems. Kinobead assays use unpurified native proteins from whole-cell lysates, which may exist in alternative forms or activation states compared to the purified recombinant proteins used in KinaseSeeker assay. Previous work has shown that *Pf*GSK3β is not essential for survival of ABS parasites, but that disruption leads to decreased growth rates and growth defects (**Supplementary Figure S4F**).^39^ Nevertheless, we tested CHIR-124 against two inducible knockdown/knockout lines of *Pf*GSK3β. A glmS-ribozyme-based inducible knockdown line (GSK3β-GFP-glmS)^39^ was generated by endogenous C-terminal tagging using selection-linked-integration (SLI),^40^ thereby introducing a glmS ribozyme sequence upstream of the 3’ untranslated region of the target gene.^41^ GlmS-mediated mRNA-destabilization was induced by treatment with 2.5 mM glucosamine (GlcN) as described previously.^42^ A DiCre-based inducible knockout line (GSK3β-DiCre) was generated by SLI^40^ using a N-terminal homology region of the target gene followed by a recodonized, loxP-flanked sequence of the C-terminus containing the kinase domain. Excision of the kinase domain coding sequence was induced by rapalog-mediated dimerization of episomally expressed DiCre as described previously.^43^ We noted no significant difference in drug sensitivity upon knockdown or knockout of *Pf*GSK3β, indicating that *Pf*GSK3β is unlikely to contribute to the mechanism of action (MoA) of CHIR-124 (**Supplementary Figure S4E-F)**. Owing to the lack of vulnerability associated with *Pf*GSK3β as a target, this result was not surprising.

### CHIR-124 Inhibits the Formation of Intracellular Hemozoin

Given the structural features of CHIR-124 that resemble hemozoin formation inhibitors (i.e. planar heterocyclic rings with basic centers,^44^ exemplified by the clinical antimalarials chloroquine (CQ) and pyronaridine **Supplementary Figure S6A**), the compound was also tested for its capacity to inhibit the formation of β-hematin (synthetic hemozoin) in an extracellular biomimetic detergent-mediated assay.^45^ Intriguingly, the IC_50_ of CHIR-124 in this assay was comparable to that of the positive control antimalarial, CQ, a validated inhibitor of hemozoin formation^46^ (**Figure 2B**), suggesting that CHIR-124 optimally interacts with free heme or crystalline β-hematin under biomimetic conditions to prevent further crystal growth. This finding prompted further whole-cell studies to better understand the effect that CHIR-124 has on the hemoglobin degradation process in the parasite. We therefore used an intracellular heme fractionation assay to probe the levels of hemoglobin, free heme and hemozoin in a dose-dependent manner when exposed to CHIR-124. While levels of hemoglobin uptake remained constant, an increase in free heme and decrease in hemozoin were observed over the range of 0.5–3×IC_50_ of CHIR-124 (**Figure 2C** and **Supplementary Figure S6B-C**). Furthermore, the increase in free heme correlated with parasite growth inhibition (**Figure 2D**). This was a notable finding given that we have also shown that *Pf*Ark1 is an efficacious target for this compound in ABS parasites. We have previously shown one other compound, TAE684, that can act as a dual *Pf*Ark1 and hemozoin formation inhibitor^37^ and, therefore, CHIR-124 expands on this, establishing an initial set of phenotypically-validated dual hemozoin formation and kinase inhibitors.

An analogous hypothesis, that the molecular structure of CHIR-124 might facilitate chelation of the compound to metal ions such as Fe(II) and Fe(III) was also tested; however, a weak capacity of CHIR-124 to bind to physiologically-relevant metal ions suggested that disruption of metal homeostasis within the parasite was unlikely to contribute to its mode of action (**Supplementary Figure S7**). We additionally explored the phenotypic effect on the growth of *Pf*NF54 parasites by combining CHIR-124 with a primary *Pf*Ark1 or hemozoin formation inhibitor. Here, we employed the Ark1 inhibitor hesperadin^37, 47^ and the hemozoin formation inhibitor chloroquine.^46^ Using this analysis, sum of the fractional IC_50_ (ΣFIC_50_) values significantly greater than 1 indicate antagonism, those between approximately 0.8 to 1.4 indicate an additive effect, while those below 0.8 indicate synergy. Fixed ratio isobologram analysis revealed antagonistic interactions for both combinations, with ΣFIC_50_ values for CHIR-124 and hesperadin or CHIR-124 and chloroquine of 1.5 and 1.7, respectively (**Figures 2E and 2F**). This profile is similar to the effect of the dual inhibitor TAE684 with CQ and hesperadin^37^, and the antagonism observed for CHIR-124 suggests that CHIR-124 interacts within the same biological pathways as both hesperadin and chloroquine, which further supports its dual modes of action.^37, 47^

### CHIR-124 Affects Early Trophozoites and Prevents Proliferation from the Schizont Phase

Considering the polypharmacology displayed by CHIR-124, we sought to further probe the contribution of these different mechanistic pathways via morphological experiments. First, we treated tightly synchronized *Pf*NF54 parasites with 3×IC_50_ of CHIR-124 at four different stages (ring, late-ring, trophozoite or schizont) described by hours post invasion (hpi) and monitored parasitemia and parasite morphology under drug pressure for every subsequent 12 h. CHIR-124 predominantly affected early trophozoites, whereby pyknotic parasites, unable to form healthy mature trophozoites, were visible within 12–24 h post treatment (hpt) of rings and early trophozoites (**Figure 3A**). Hemozoin crystal formation was also visibly affected and the nuclei in these parasites were unable to divide to form schizonts, with the nuclear content per cell (n) dramatically diminished in treated parasites compared to the untreated controls over 72 h (**Figure 3B**). This effect is especially striking at time points 24 h and 72 h (30–34 hpi and 78–82 hpi, respectively), which represent the schizont stages of untreated parasites. Analysis of the cell distribution via staining of parasite DNA with SYBR Green I further confirmed that CHIR-124-treated parasites were stalled in the early trophozoite phase (12 h) relative to the untreated population, which showed a shift to the right (**Figure 3B**). However, CHIR-124 had little morphological effect on parasites already in the mature trophozoite phase (36 hpi), with hemozoin crystals visible (**Figure 3A**). These parasites were able to enter schizogony, albeit with aberrant structures and fewer nuclei relative to the untreated control. Finally, parasites treated at 48 hpi that had already entered schizogony, went on to form some new rings but at lower parasitemias than the untreated control. This profile is consistent with the dual inhibition of a kinase as well as inhibiting hemozoin formation, as seen with TAE684, but not for a sole *Pf*Ark1 inhibitor such as hesperadin.^37^

**Figure 3.**
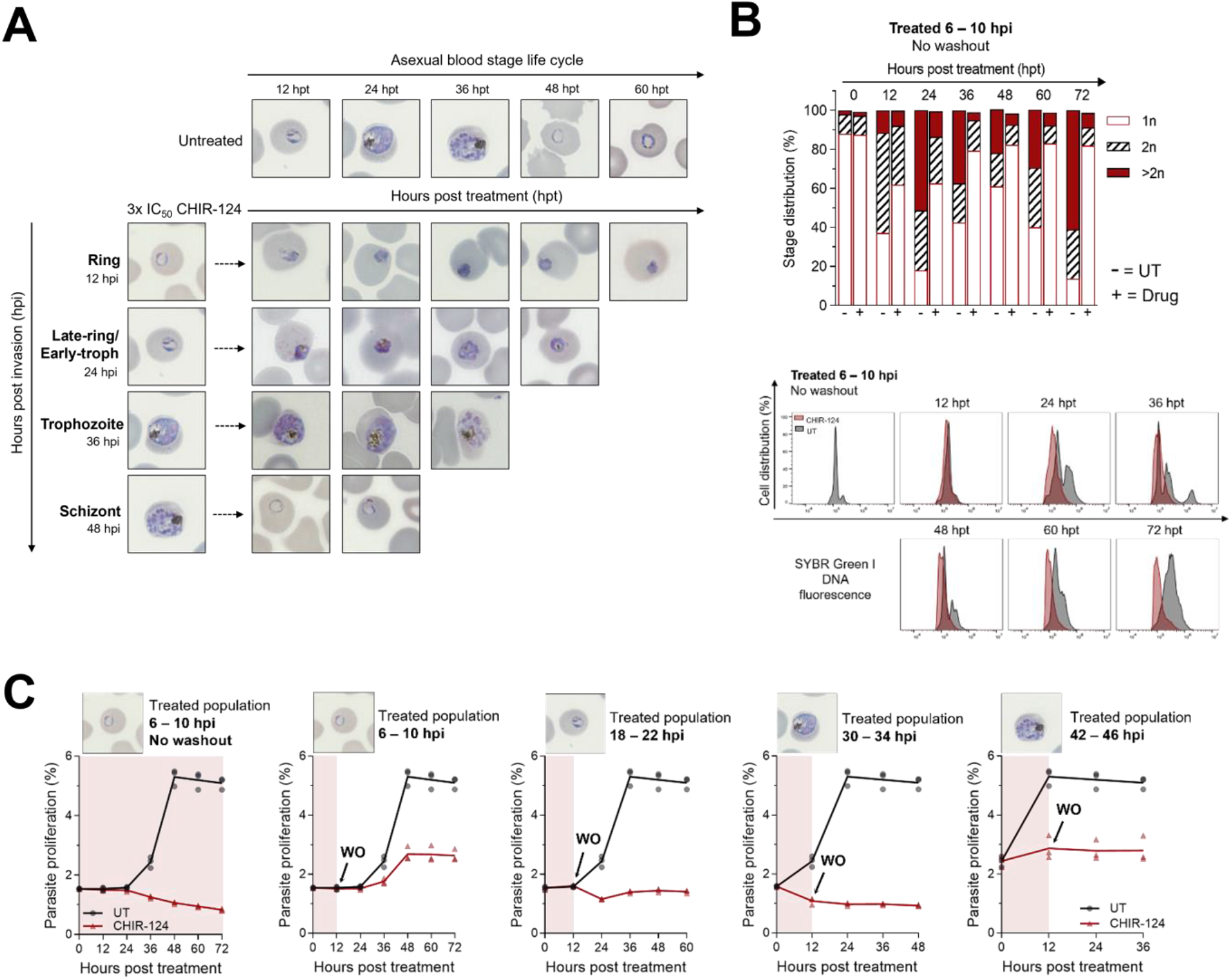
The morphology of ABS parasites treated with CHIR-124 at 3×IC_50_ vs untreated (UT) at various time-points indicated by hours post invasion (hpi) and samples at specific hours post treatment (hpt) where (**A**) shows images of Giemsa-stained blood smears, (**B**) shows the percentage stage distribution of the ABS parasites where schizonts are represented by number of nuclei per cell (n) as >2n, and (**C**) the growth of parasites when distinct synchronized populations are treated with CHIR-124 for 12 h.

To quantify the reduction of new rings after treatment of schizonts with CHIR-124 as well as to determine if the effect of CHIR-124 is reversible, we then performed similar experiments with a washout of CHIR-124 after 12 h and monitored the parasitemia in 12 h intervals via flow cytometry (**Figure 3C**). Regardless of the starting point of treatment (measured at hpi), CHIR-124-treated parasites were unable to proliferate relative to the untreated control, despite the removal of drug pressure. These findings indicate that, while CHIR-124 has the most significant morphological effect during the trophozoite phase due to its inhibition of hemozoin formation, it can arrest the growth of parasites treated during the schizont phase. In the latter, hemozoin formation is no longer relevant.^48^ Instead, this observation supports the inhibition of *Pf*Ark1, which is crucial for cell division and parasite proliferation. *Pf*Ark1 is one of the Aurora-related (serine/threonine) kinases that regulates microtubule spindle dynamics during the parasite’s mitotic process, leading to defects in nuclear segregation and ultimately daughter cell fromation.^37, 49^ Indeed, *Pf*Ark1 has been shown to be essential for erythrocytic schizogony and is localized to spindle pole bodies.^50, 51^ The abnormal schizont morphology and lower-yield nuclear division correlates with that reported for *Plasmodium* parasites treated with the *Pf*Ark1 inhibitor hesperadin.^47^ These data provide further evidence of CHIR-124’s ability to inhibit both hemozoin formation and the *Plasmodium* kinase, *Pf*Ark1. Interestingly, although *Pf*CDPK1 is not essential for asexual blood stage survival, one study suggests that its inhibition affects RBC invasion.^52^ Therefore, inhibition of *Pf*CDPK1, which was also competed off the Kinobeads may contribute to the reduced parasitemia after treatment of schizonts with CHIR-124.

### The Risk of Resistance is Low for Parasites Treated with CHIR-124

One potential advantage of a compound that targets multiple pathways in the parasite is the lowered risk of resistance-conferring mutations arising simultaneously. To further probe the frequency of resistance for CHIR-124, 10^9^ *Pf*Dd2 parasites were pressured with non-lethal doses of the compound (3×IC_50_) until complete parasite clearance from the culture as visualized by microscopic examination from Giemsa-stained blood smears. The drug pressure was removed until parasites recrudesced (49 days after initializing the resistance selection). These parasites, however, showed no resistance phenotype (**Supplementary Figure S8**) and were therefore re-exposed to drug pressure at 5×IC_50_ until cleared, in attempt to select for higher-grade resistance. The culture was continued at 3×IC_50_ for 60 days. At this point, no recrudescence had been observed, indicating that the minimum inoculum of resistance (MIR) under the *in vitro* experimental conditions for CHIR-124 is log(MIR) >9 against *Pf*Dd2. The lack of resistant mutants demonstrated a low level of risk for generating resistance against CHIR-124, and supported the observation of polypharmacology.^53^

## Discussion

The present study highlights the potential of target repurposing strategies in antimalarial discovery by demonstrating that the human Chk1 inhibitor CHIR-124 exerts potent activity against *P. falciparum* through a dual mechanism involving *Pf*Ark1 inhibition and the disruption of hemozoin formation. While the discovery of new inhibitors of hemozoin formation is not generally a priority for MMV, given that the hemozoin formation pathway is restricted to the asexual blood stage (i.e. TCP1), this class often shows advantages such as excellent selectivity for the parasite and associations with lower risks of resistance. Additionally, a significant percentage of potent analogs from phenotypic screens are likely to have some activity in β-hematin formation assays.^54, 55^ Therefore, continued investigation of the heme detoxification pathway remains relevant and beneficial for antimalarial drug development.^48^

Interestingly, the identification of CHIR-124 originated from a screen to discover *Plasmodium* kinase inhibitors, as opposed to hemozoin formation inhibitors. However, this is not the first time a chemical series has presented the combination of inhibitors of a kinase and hemozoin formation.^37, 56-58^ The reason behind this phenomenon comes from the overlapping molecular structural features, including multiple heteroaromatic rings and basic nitrogen atoms, present in both hemozoin formation inhibitors and kinase inhibitors.^56^ Notably, in a previous study of 2,8-disubstituted-1,5-naphthyridines, the MoA switched from kinase inhibition to hemozoin formation inhibition between chemical derivatives.^57^ In another study of a type II human kinase inhibitor, hemozoin formation was disrupted and biochemical inhibition of recombinant *P. falciparum* protein kinase 6 (*Pf*PK6) was shown; however no IC_50_ shift was observed against a knockdown of *Pf*PK6, suggesting the inhibition hemozoin formation may be the driver of whole-cell activity.^58^

On the other hand, the present work provides whole-cell phenotypic evidence via the use of knockdowns and cell fractionation that CHIR-124 combines kinase and hemozoin formation inhibition in the same molecule, with each mechanism contributing to antiplasmodial activity. Furthermore, CHIR-124 acts on both trophozoites and schizonts, presumably via these independent biological pathways. This polypharmacological profile is particularly significant given the increasing prevalence of resistance to frontline therapies such as artemisinin and partner drugs in artemisinin-based combination therapies (ACTs). By simultaneously disrupting heme detoxification and inhibiting a parasite kinase essential for cell cycle regulation, the likelihood of resistance development is reduced, as parasites would need to acquire mutations in multiple pathways to overcome its activity. Similar approaches targeting multiple parasite processes have been proposed as a strategy to prolong drug efficacy and delay resistance emergence.^59^

The identification of *Pf*Ark1 as a relevant target aligns with growing evidence that *Plasmodium* kinases represent a promising yet underexplored class of drug targets. Recent structure–activity relationship studies on Aurora-related kinases and other parasite kinases have underscored their essential roles in parasite proliferation and validated them as tractable targets for medicinal chemistry optimization.^59^ The availability of human Aurora kinase structures that can be used in combination with a homology model to guide the design of selective inhibitors, as well as the key differences in the ATP binding sites between the human and *Plasmodium* Ark orthologues, also makes *Pf*Ark1 suitable and attractive for structure-guided drug development.^37^ Notably, the *Pf*Ark1 inhibitor, hesperadin, shows a much greater affinity for *Pf*Ark1 than CHIR-124 in biochemical assays, but the inverse was found in *Pf*Ark1 cKD experiments with CHIR-124 demonstrating a larger degree of sensitization. This is not unexpected since differences in IC_50_ shifts against whole-cell cKDs are not typically correlated with variation in extracellular target activity, especially when polypharmacology is involved, as observed in the case of TAE684. However, interestingly, the respective activity against the recombinant form of the protein is within 2-fold of the corresponding ABS whole-cell potency for hesperadin, TAE684 and CHIR-124.^37^

The inhibition of hemozoin formation by CHIR-124 provides an additional layer of activity reminiscent of quinoline antimalarials, but with a distinct chemical scaffold, potentially circumventing cross-resistance issues. Moreover, the moderate activity observed against liver and gametocyte stages further shows the benefits of the additional kinase inhibition within the same molecule and suggests that future CHIR-124 derivatives could contribute to transmission-blocking strategies, an increasingly important goal in malaria elimination efforts.^12^ Taken together, these findings support the feasibility of identifying dual-action inhibitors that combine kinase inhibition with disruption of heme detoxification. Such compounds may represent a new generation of antimalarials with reduced resistance risks and multistage activity. Future work should focus on optimizing the *Pf*Ark1 activity, parasite selectivity and pharmacokinetic properties of CHIR-124 analogs, assessing *in vivo* efficacy, and exploring synergistic combinations with existing antimalarials to maximize therapeutic potential. Ultimately, the development of such multitargeted therapies through medicinal chemistry optimization could significantly advance malaria control and eradication efforts.

## Supporting Information

The authors have cited additional references within the Supplementary Information.^[60-96]^

Supplementary Figures S1 – S7, Supplementary Tables S1 – S4, protocols for Kinobead screening, *Plasmodium falciparum* asexual blood stage parasitology methods, gametocyte and liver stage assays, cytotoxicity assays, stage specificity assays, conditional knockdown assays, β-hematin and hemozoin formation inhibition assays, metal chelation assays and biochemical assays.

## Data availability statement

The data supporting this article have been included as part of the Supplementary Information.

## Acknowledgements

South African Medical Research Council extramural unit funding to K.C. Medicines for Malaria Venture (MMV) African Challenge Grant RD-17-0047 awarded to D.C. and L-M.B and Project RD-19-0001 to L-M.B. Future Leaders−African Independent Research (FLAIR) Fellowship Programme, a partnership between the African Academy of Sciences and the Royal Society funded by the UK Government’s Global Challenges Research to L.B.C. The National Institute of Allergy and Infectious Diseases of the National Institutes of Health to K.J.W. (R01AI143521), D.A.F. (R01AI185559) and K.C. (R01AI152092). E.A.W. is a member of the Malaria Drug Accelerator and is supported by a grant from the Bill and Melinda Gates Foundation (OPP1054480). The Department of Science and Innovation and the National Research Foundation South African Research Chair (SARChI) in Sustainable Malaria Control to L-M.B. (UID 84627). K.C. is the Neville Isdell Chair in African-centric Drug Discovery and Development and thanks Neville Isdell for generously funding the Chair.

## Author Contributions

The manuscript was prepared by K.J.W. with contributions from all co-authors. β-hematin and metal chelation studies were conducted by J.G.W. Cell fractionation and combination studies were performed or analyzed by L.F.G. and K.J.W. Morphological studies were carried out by H.L. and L-M. B. Asexual blood stage parasite activity assays were performed by D.T., J.L.B. and D.A.F. Gametocyte assays were conducted by D.C., J.R, M.vdW and L-M.B. Liver-stage assays were done by M.L.S, J.L.S-N. and E.A.W. Kinobead experiments were conducted by S.G-D, M.J.L-M and F-J.G. Conditional knockdown experiments were performed and analyzed by L.C.G, C.F.P., J.C.N, C.S., S.W., T.S.V., A.A. and T-W.G. Barcoded mutant screening was conducted by G.G., R.C. and M.L. Resistance selections were carried out by K.J.W. The project was conceived and supervised by K.J.W., L.B.C and K.C.

## Entry for the Table of Contents

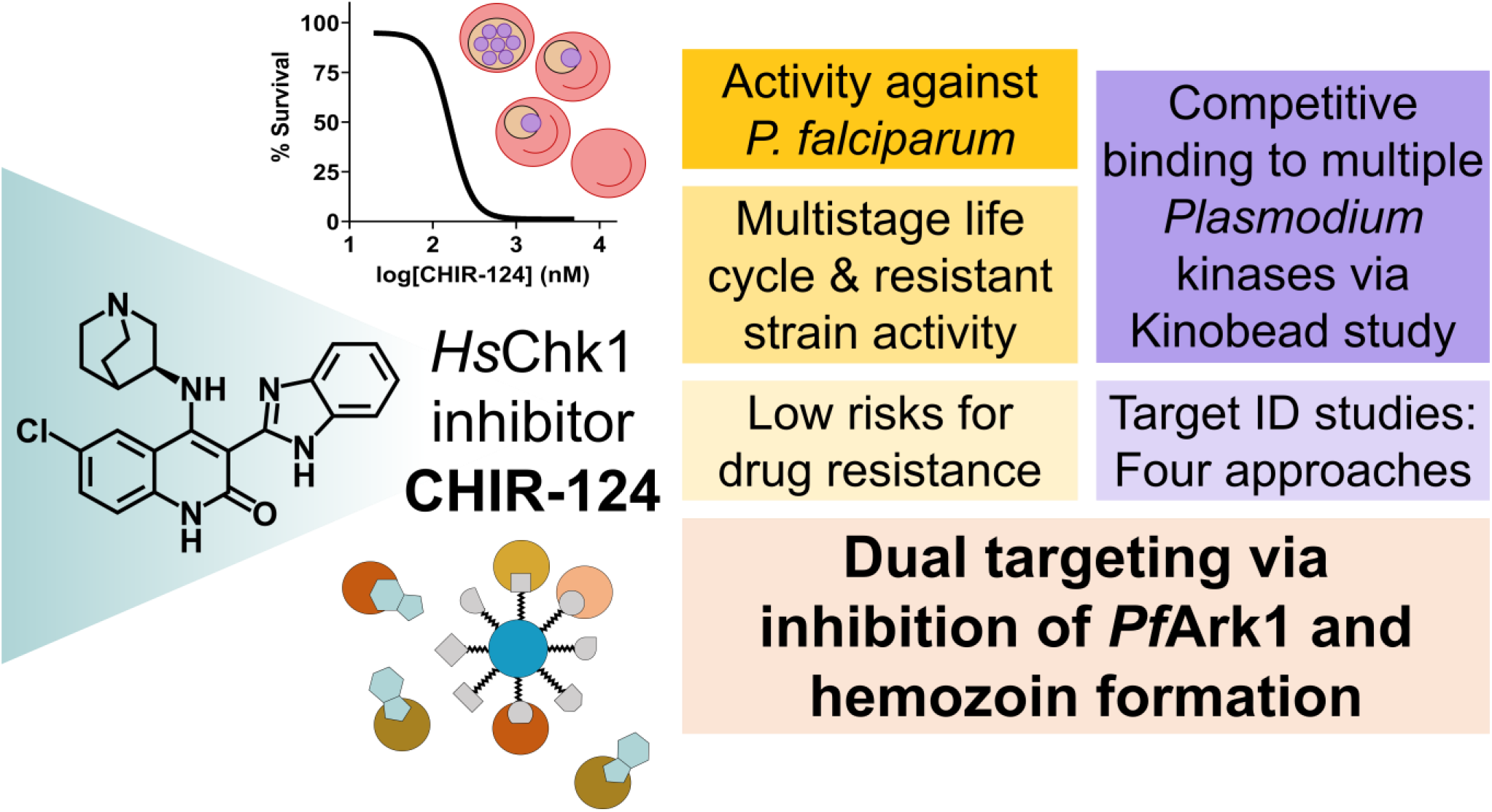

