## Supplementary Information for "The Human Chk1 Inhibitor CHIR-124 Shows Multistage Activity Against the Human Malaria Parasite *Plasmodium falciparum* via Polypharmacological Inhibition of *Pf*Ark1 and Hemozoin Formation"

### CONTENTS


#### 1. Supplementary Figures

Supplementary Figure S1 – Competitive inhibition data from Kinobeads assay
Supplementary Figure S2 – Cross-resistance with Southeast Asian lines
Supplementary Figure S3 – Cross-resistance with DNA-barcoded resistant mutant lines
Supplementary Figure S4 – Conditional knockdown (cKD) studies
Supplementary Figure S5 – Biochemical *Plasmodium* kinase activities
Supplementary Figure S6 – Cellular heme fractionation data
Supplementary Figure S7 – Metal chelation studies
Supplementary Figure S8 – Resistance selection profiling

#### 2. Supplementary Tables

Supplementary Table S1 – Kinobead MS/MS Data
Supplementary Table S2 – $IC_{50}$ data for asexual blood stage activities
Supplementary Table S3 – Liver stage activity and cytotoxicity data
Supplementary Table S4 – List of barcoded lines in the AReBar cross-resistance screening pool
Supplementary Table S5 – Conditional knockdown (cKD) data
Supplementary Table S6 – List of oligonucleotides for donor vector construction

#### 3. Methods

3.1 Kinobead screening

3.2 *Plasmodium falciparum* parasitology methods
- 3.2.1 Parasite culture
- 3.2.2 Antiplasmodial activity assessments
- 3.2.3 Stage specificity assay
- 3.2.4 Stage-specific gametocyte production and luciferase assay
- 3.2.5 Liver stage assay
- 3.2.6 Liver stage cytotoxicity
- 3.2.7 Conditional knockdown assays
- 3.2.8 Cross-resistance profiling using the Antimalarial Resistome Barcode sequencing assay
- 3.2.9 Combination studies via fixed-ratio isobolograms
- 3.2.10 Rate- and stage-specific morphological evaluations and inhibitor effect reversibility

3.3 β-hematin and hemozoin formation inhibition assays and metal chelation studies
- 3.3.1 NP-40 based extracellular β-hematin inhibition assay
- 3.3.2 Cellular heme fractionation assay
- 3.3.3 Metal chelation studies

3.4 Biochemical assays
- 3.4.1 KinaseSeeker™ assay

### 1. Supplementary Figures

#### Fig. S1
**A**

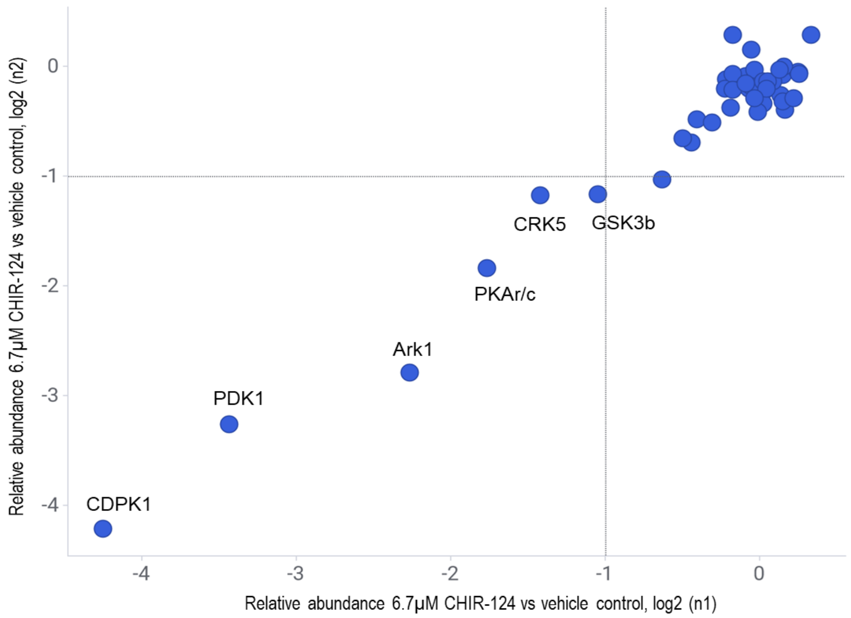


**B**

| Gene name | UniProt ID | Protein description | Former name | $IC_{50}$ (µM) n=2 | $K_d^{app}$ (µM) |
|---|---|---|---|---|---|
| PF3D7_0217500 | P62344 | CDPK1, Calcium-dependent protein kinase 1 | PFB0815W | 0.0823* | 0.04 |
| | | | | 0.0823* | 0.04 |
| PF3D7_1121900 | Q8IIE7 | PDK1, Serine/threonine protein kinase, putative | PF11_0227 | 0.21 | 0.12 |
| | | | | 0.15 | 0.08 |
| PF3D7_0605300 | C6KSQ1 | Ark1, Serine/threonine protein kinase | PFF0260W | 0.32 | 0.18 |
| | | | | 0.28 | 0.16 |
| PF3D7_1223100 | Q7KQK0 | PKAr, CAMP-dependent protein kinase regulatory subunit, putative | PFL1110C | 0.83 | 0.64 |
| | | | | 0.87 | 0.61 |
| PF3D7_0934800 | Q7KQA0 | PKAc, cAMP-dependent protein kinase catalytic subunit | PFI1685W | 0.88 | 0.63 |
| | | | | 0.85 | 0.59 |
| PF3D7_0615500 | C6KSZ6 | CRK5, Cdc2-related protein kinase 5 | PFF0750W | 1.7 | 1.04 |
| | | | | 2.9 | 1.89 |
| PF3D7_0312400 | O77344 | GSK3beta, Glycogen synthase kinase 3 | PFC0525C | 2.8 | 1.23 |
| | | | | 3.1 | 1.32 |
| PF3D7_0805700 | C0H4R8 | FIKK8, Serine/threonine protein kinase, FIKK family | MAL8P1.203 | 8.2 | 7.87 |
| | | | | 3.5 | 2.97 |

*0.0823 µM was the lowest compound concentration used.

**Supplementary Figure S1.** Kinobeads profiling of CHIR-124 was conducted using a dose-dependent competition assay (starting at 20 µM, followed by serial 1:3 dilutions). (**A**) Schematic representation of two independent experiments. (**B**) Summary table detailing $IC_{50}$ values and apparent $K_d$ values, which reflect the binding affinity of the protein to the compound immobilized on the beads. Data are reported for the seven competed kinases, including the regulatory subunit of PKAr/c.

### Fig. S2

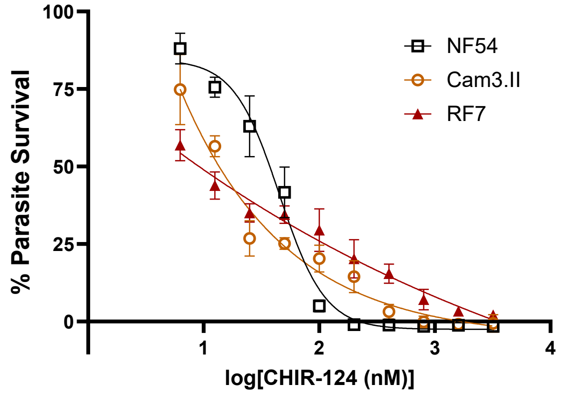


**Supplementary Figure S2.** Cross resistance with Southeast Asian strains. CHIR-124 was profiled against the *P. falciparum* lines Cam3.II (resistant to chloroquine and artemisinin) and RF7 (resistant to artemisinin and piperaquine) and displayed more shallow dose response curves.

### Fig. S3

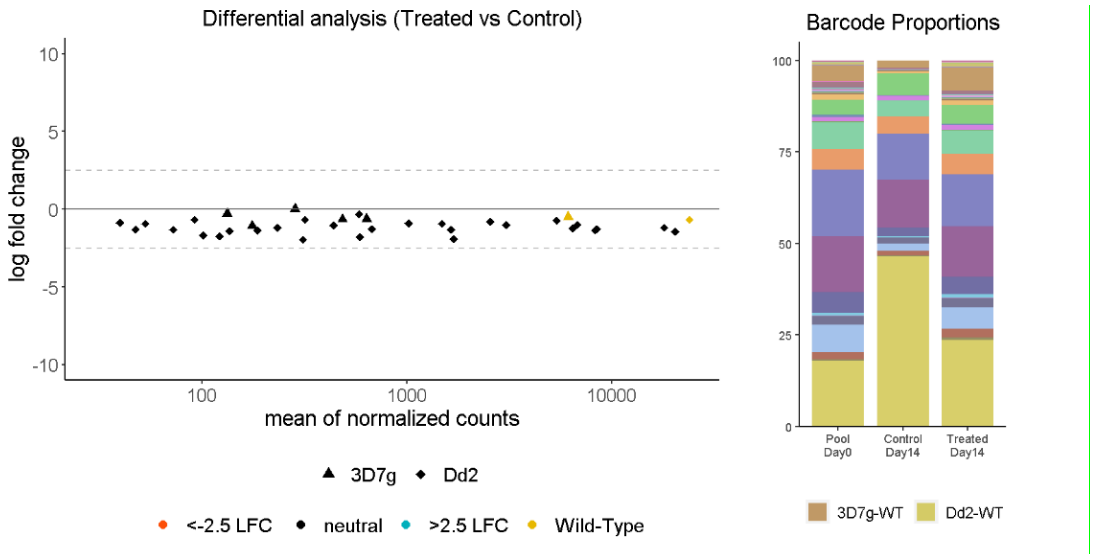


**Supplementary Figure S3.** Cross resistance profiling using the Antimalarial Resistome Barcode sequencing (AReBar) assay. A pool of 45 DNA-barcoded resistant mutant lines (**Supplementary Table S4**) were exposed to 3×$IC_{50}$ concentrations of CHIR-124. No mutant strain enrichment was observed after 14 days. *Left panel*, differential analysis comparing CHIR-124-treated parasites versus untreated control pool, with the log2 fold change (LFC) of individual barcoded lines (y-axis) and barcode counts (x-axis) shown. Dotted lines indicate ±2.5 LFC relative to untreated control lines, and symbols indicate strain background of mutant lines. *Right panel*, barcode proportions of each line at day 0, and day 14 for the untreated control and CHIR-124-treated pool.

**Fig. S4**

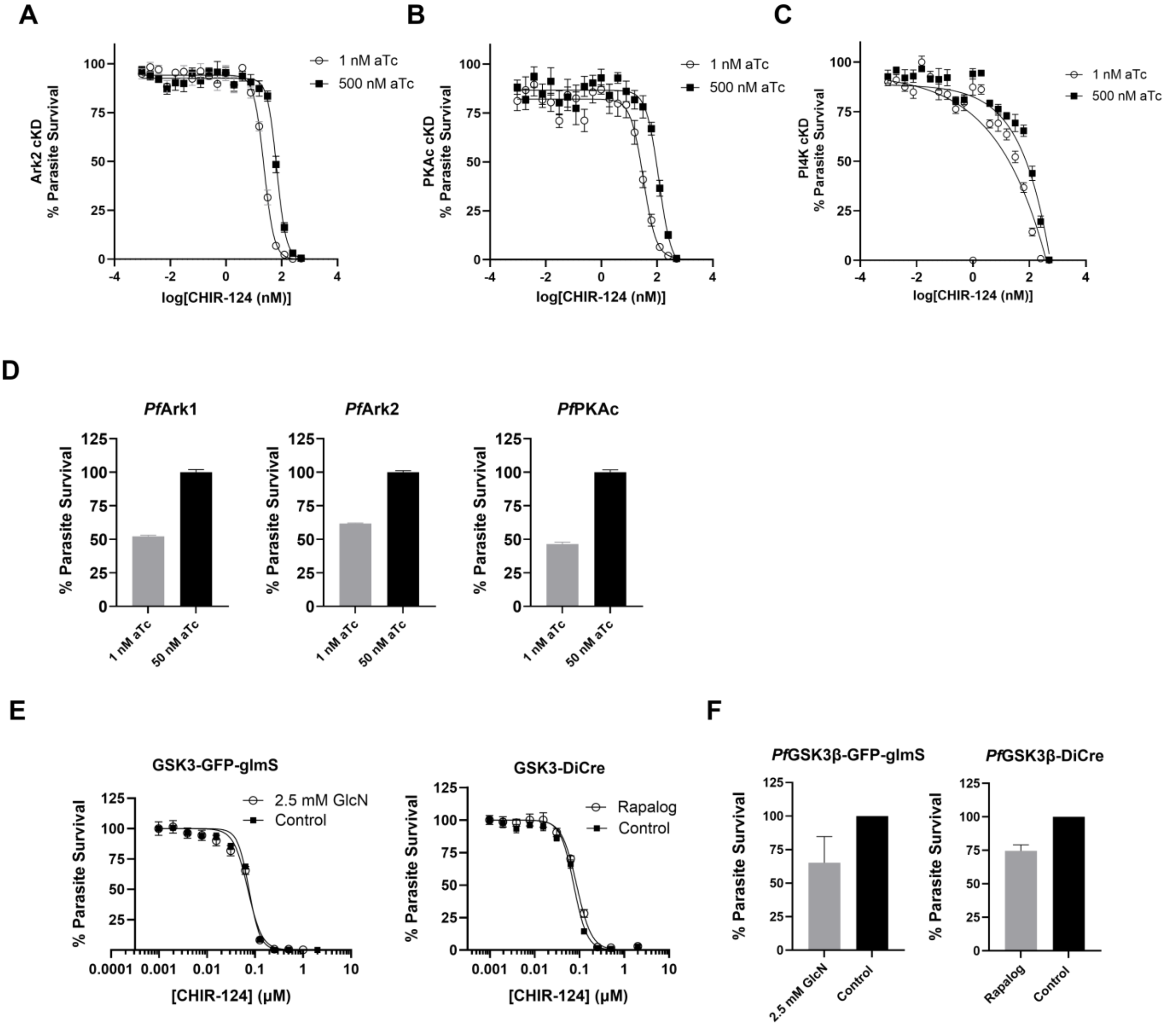


**Supplementary Figure S4.** Conditional knockdown (cKD) studies with CHIR-124. TetR-based anhydrotetracycline (aTc)-controlled conditional knockdown (cKD) studies with the (**A**) *Pf*Ark2 cKD line, (**B**) *Pf*PKAc cKD line and (**C**) *Pf*PI4K cKD line whereby expression of the kinase was regulated by the concentration of aTc on the culture during the 72 h assay. The *Pf*PI4K cKD represents the negative control demonstrating that the small (2–3 fold) curve shift to the left for the culture under 1 nM aTc (grey), relative to the 500 nM aTc line where the kinase was fully expressed, indicates a negligible interaction between CHIR-124 and the specified kinase targets. Data represents mean ± SEM where N,n = 3,3. (**D**) Parasite viability for untreated cKD lines used in this study represented by 72 h growth assays in presence of high (50 nM, WT condition) and low (1 nM, knockdown) aTc. Data represents mean ± SEM where N,n = 3,3. Viability data for the *Pf*PI4K cKD has been previously reported.[60] (**E**) CHIR-124 tested against inducible lines *Pf*GSK3β-GFP-glmS and *Pf*GSK3β-DiCre comparing 2.5 mM glucosamine (GlcN) or 250 nM rapalog versus untreated control respectively. No difference in drug sensitivity was observed in a 96 h proliferation assay indicating that *Pf*GSK3β is unlikely to contribute to the mode of action of CHIR-124. Data represents mean ± SEM where N,n = 3,2). (**F**) Parasite viability for untreated *Pf*GSK3β-GFP-glmS or *Pf*GSK3β-DiCre used in this study represented by 96 h growth assays normalized to the control (100% survival) in the presence of 2.5 mM GlcN or rapalog, respectively. Data represents mean ± SEM where N = 3-4.

### Fig. S5

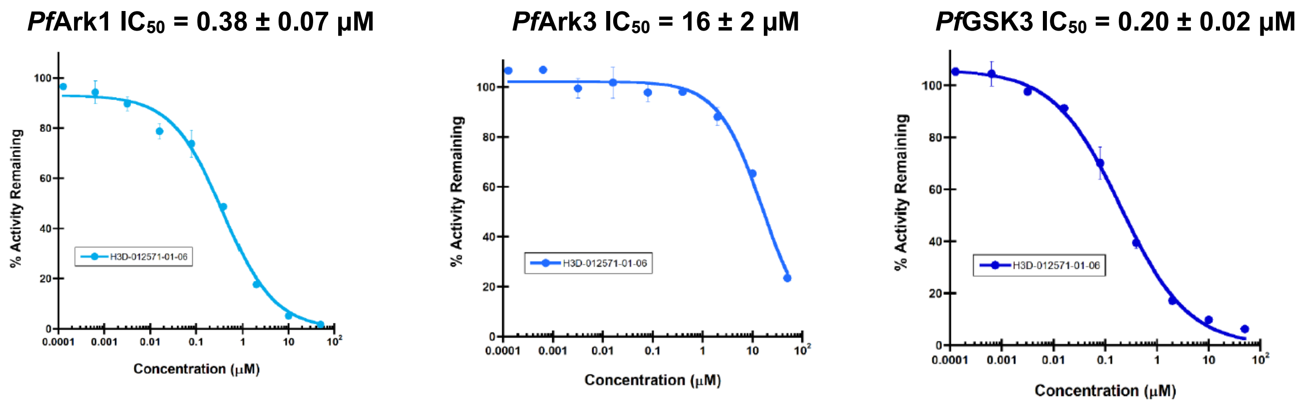


**Supplementary Figure S5.** *Plasmodium* kinase biochemical activities against *Pf*Ark1, *Pf*Ark3 and *Pf*GSK3β. Mean ± SD shown above their dose-response curves with error bars representing SD from screening against the specified kinase in duplicate.

### Fig. S6

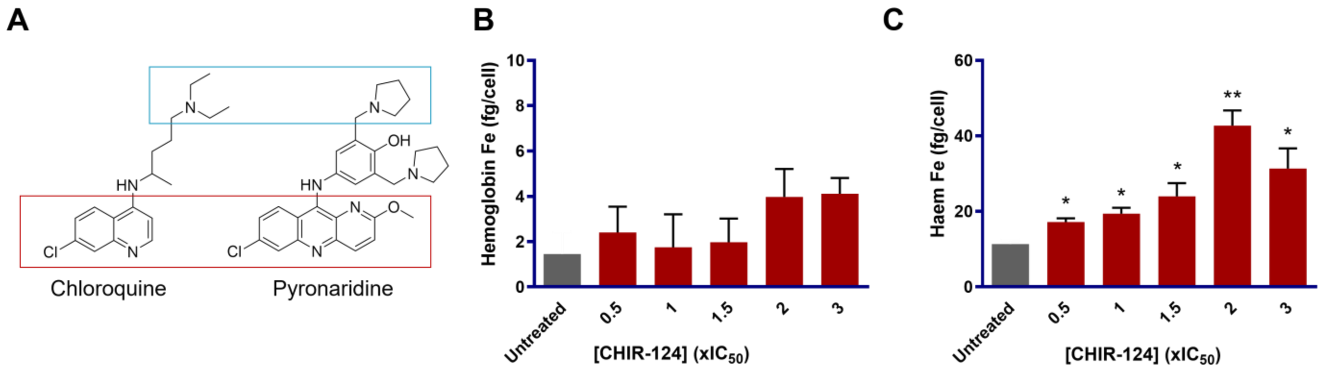


**Supplementary Figure S6:** (**A**) Validated clinical hemozoin formation inhibitors, chloroquine and pyronaridine, highlighting the typical pharmacophore for this class of inhibitor which contains planar heteroaromatics (red box) and basic centers (blue box). Cellular heme fractionation assays with multiples of the CHIR-124 $IC_{50}$ showing heme Fe (fg/cell) levels from the (**B**) free hemoglobin fraction, and (**C**) absolute free heme fraction.

Fig. S7

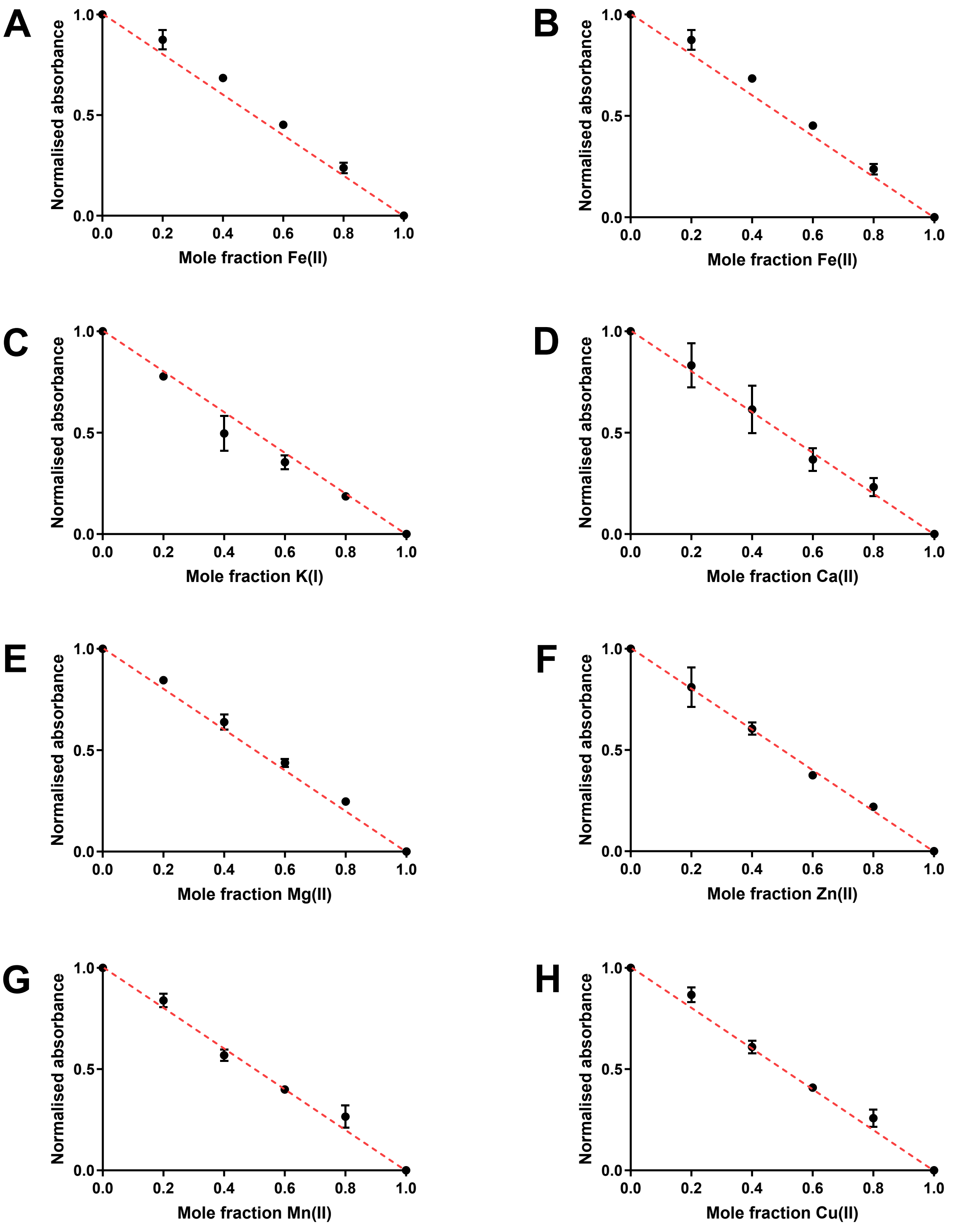


**Supplementary Figure S7.** Metal chelation studies via spectrophotometric titrations of CHIR-124 with (**A**) Fe(**II**), (**B**) Fe(**III**), (**C**) K(**I**), (**D**) Ca(**II**), (**E**) Mg(**II**), (**F**) Zn(**II**), (**G**) Mn(**II**) and (**H**) Cu(**II**) showing, in all cases, a very weak capacity for chelation of these metal ions with CHIR-124. Dashed red lines indicate Beer-Lambert law dilutions curves; deviation from that line indicates likely chelation. Error bars represent technical duplicates of each experiment.

**Fig. S8**

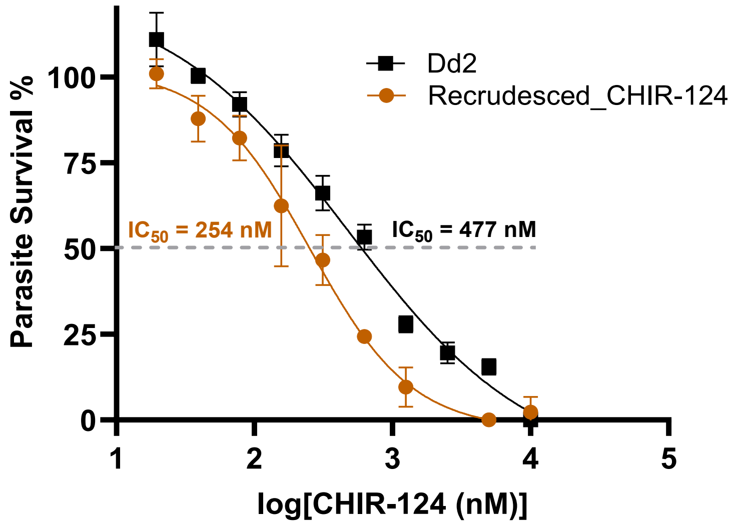


**Supplementary Figure S8:** Resistance selection profiling. Parasites recrudesced while the flask was being cultured without CHIR-124 pressure on day 49. The parasites were subjected to a pLDH assay on one occasion whereupon they were found to be approximately 2-fold more sensitive to CHIR-124 and subsequently put back on CHIR-124 pressure at 5×$IC_{50}$ without further profiling. Data points and error bars represent mean ± SD from one biological repeat.

### 2. Supplementary Tables

**Supplementary Table S1.** Kinobeads MS/MS Data**.** *See Excel file.* MS/MS data for profiling of CHIR-124 (GSK3210608, CAS No. 405168-58-3) on Kinobeads in *P. falciparum* extracts: 10 samples were analyzed in parallel (TMT 10-plex) to generate values for the affinity of the beads to the bound proteins ("depletion" values, 4 samples) and to generate $IC_{50}$ values (6 samples) in a single experiment. Samples 1 and 2 represent the vehicle control. Samples 3 and 4 were done in the same way, but while the beads were discarded after the first incubation step the extract was incubated with fresh beads to measure how much protein could rebind to the fresh beads (protein was depleted from the extract by first bead-binding). Apparent dissociation constants were determined by taking into account the protein depletion by the Kinobeads. Samples 5–10 were used to generate $IC_{50}$ values by adding compound over a range of concentrations (20 µM, 1:3 dilutions).

| Experiment number | Compound immobilized on beads | "Free" compound added to extract for competition | Sample description | Compound Concentration (µM) |
|---|---|---|---|---|
| X048537 | Kinobeads | GSK3210608 CHIR-124 | Vehicle control (n=1) | 0 |
| | | | Vehicle control (n=2) | 0 |
| | | | Re-binding of NBF to fresh beads (n=1) | 0 |
| | | | Re-binding of NBF to fresh beads (n=2) | 0 |
| | | | Dose response competition over 6 concentrations | 20.0 |
| | | | | 6.67 |
| | | | | 2.22 |
| | | | | 0.74 |
| | | | | 0.25 |
| | | | | 0.08 |
| X048538 | Kinobeads | GSK3210608 CHIR-124 | Vehicle control (n=1) | 0 |
| | | | Vehicle control (n=2) | 0 |
| | | | Re-binding of NBF to fresh beads (n=1) | 0 |
| | | | Re-binding of NBF to fresh beads (n=2) | 0 |
| | | | Dose response competition over 6 concentrations | 20.0 |
| | | | | 6.67 |
| | | | | 2.22 |
| | | | | 0.74 |
| | | | | 0.25 |
| | | | | 0.08 |

**Supplementary Table S2.** $IC_{50}$ data for asexual blood stage activities. Inhibition of the WT drug-sensitive (*Pf*NF54) and drug-resistant strains by CHIR-124, showing the mean $IC_{50}$ ± SEM and Hill slope calculated from N,n = 2-3,2 repeats.

| Strain | Assay detection | Mean $IC_{50}$ ± SEM (µM) | Mean Hill slope |
|---|---|---|---|
| *Pf*NF54 (WT) | pLDH | 0.14 ± 0.03 | -2.9 ± 0.4 |
| *Pf*Dd2 | pLDH | 0.44 ± 0.05 | -1.3 ± 0.5 |
| *Pf*K1 | pLDH | 0.14 ± 0.03 | -1.1 ± 0.2 |
| *Pf*NF54 (WT) | SYBR Green I | 0.033 ± 0.002 | -2.2 ± 0.2 |
| *Pf*Cam3.II | SYBR Green I | 0.016 ± 0.002 | -0.5 ± 0.3 |
| *Pf*RF7 | SYBR Green I | 0.010 ± 0.001 | -0.1 ± 0.7 |

**Supplementary Table S3.** Liver stage activity and cytotoxicity data.

| | Liver cytotoxicity HepG2 $IC_{50}$ (µM) | Liver-stage PbLuc $IC_{50}$ (µM) |
|---|---|---|
| Rep 1 | 5.4 | 0.30 |
| Rep 2 | 4.3 | 0.34 |
| Rep 3 | 3.3 | 0.30 |
| **Mean** | **4.3** | **0.31** |
| **SEM** | **0.6** | **0.01** |

**Supplementary Table S4. List of barcoded lines in the AReBar cross-resistance screening pool**. The line name, indicating strain background (*Pf*3D7 or *Pf*Dd2) and mutation are shown in column 1. Gene description, gene ID, and proportion of each line in the pool at day 0 and after 14 days of CHIR-124 treatment (3×$IC_{50}$) are shown.

| Line name | Gene Description | Gene ID | Input Day 0 (%) | CHIR-124 Day 14 (%) |
|---|---|---|---|---|
| 3D7-WT | Wild type | | 4.31 | 6.43 |
| 3D7 ABCI3 R2180P | ABC transporter I family member 1 | PF3D7_0319700 | 0 | 0.02 |
| 3D7 ACS10 M300I | Acyl CoA synthase | PF3D7_0525100 | 0.39 | 0.48 |
| 3D7 ACS11 D648Y | Acyl CoA synthase | PF3D7_1238800 | 0 | 0 |
| 3D7 ACS11 E668K | Acyl CoA synthase | PF3D7_1238800 | 0.01 | 0 |
| 3D7 ATP2 CNV2 | Phospholipid-transporting ATP2 | PF3D7_1219600 | 0.01 | 0.03 |
| 3D7 DHFR-TS G378E | Dihydrofolate reductase-thymidylate synthase | PF3D7_0417200 | 0.16 | 0.14 |
| 3D7 DHFR-TS I403L | Dihydrofolate reductase-thymidylate synthase | PF3D7_0417200 | 0.17 | 0.37 |
| 3D7 FTb A515T | Farnesyltransferase subunit beta | PF3D7_1147500 | 0.49 | 0.63 |
| 3D7 MDR2 K840N | Multidrug resistance protein 2 | PF3D7_1447900 | 0.03 | 0.02 |
| 3D7 NCR1 A1108T | Niemann-Pick type C1-related protein | PF3D7_0107500 | 0.09 | 0.15 |
| Dd2-WT | Wild type | | 18.08 | 23.58 |
| Dd2 AcAS A597V | Acetyl-CoA synthetase | PF3D7_0627800 | 0.12 | 0.07 |
| Dd2 ATP4 G358S | Non-SERCA-type Ca2+ -transporting P-ATPase (ATP4) | PF3D7_1211900 | 1.62 | 0.91 |
| Dd2 ATP4 A353E CARL I1139K | ATP4+CARL | PF3D7_1211900 | 0.09 | 0.06 |
| Dd2 CARL I1139K | Cyclic amine resistance locus (CARL) | PF3D7_0321900 | 0.16 | 0.13 |
| Dd2 CARL L1073Q | Cyclic amine resistance locus (CARL) | PF3D7_0321900 | 0.07 | 0.09 |

| | | | | |
|---|---|---|---|---|
| Dd2 CARL V1103L | Cyclic amine resistance locus (CARL) | PF3D7_0321900 | 0.13 | 0.09 |
| Dd2 CPSF Y408S (edit) | Cleavage and polyadenylation specific factor | PF3D7_1438500 | 0.01 | 0.01 |
| Dd2 CPSF Y408S (sel) | Cleavage and polyadenylation specific factor | PF3D7_0709000 | 0 | 0 |
| Dd2 CRT M343L | Chloroquine resistance transporter | PF3D7_1250200 | 0.59 | 0.5 |
| Dd2 CSC1 L800P | CSC1-like protein putative | PF3D7_MIT02300 | 0.25 | 0.31 |
| Dd2 cytBC1 G33V | Cytochrome b | PF3D7_MIT02300 | 0.21 | 0.17 |
| Dd2 cytBC1 V284L | Cytochrome b | PF3D7_0417200 | 0.32 | 0.36 |
| Dd2 DHFR-TS S216R | Dihydrofolate reductase-thymidylate synthase | PF3D7_0603300 | 1.43 | 1.2 |
| Dd2 DHODH C276Y | Dihydroorotate dehydrogenase | PF3D7_0603300 | 4.01 | 5.14 |
| Dd2 DHODH F227I | Dihydroorotate dehydrogenase | PF3D7_0603300 | 0.02 | 0.01 |
| Dd2 DHODH I263F | Dihydroorotate dehydrogenase | PF3D7_1451100 | 0.02 | 0.01 |
| Dd2 DHODH L531F | Dihydroorotate dehydrogenase | PF3D7_1451100 | 0.05 | 0.04 |
| Dd2-pol-delta D308A-E310A | DNA polymerase delta | PF3D7_1017000 | 0.56 | 0.34 |
| Dd2 eEF2 L755F | Elongation factor 2 | PF3D7_1128400 | 1.18 | 1.3 |
| Dd2 eEF2 Y186N | Elongation factor 2 | PF3D7_1332900 | 0.03 | 0.03 |
| Dd2 GGPPS S228T | Geranylgeranyl diphosphate synthase | PF3D7_1332900 | 0.31 | 0.16 |
| Dd2 IleRS E180D | Ile-tRNA synthetase | PF3D7_1332900 | 7.15 | 6.25 |
| Dd2 IleRS L810F | Ile-tRNA synthetase | PF3D7_1343700 | 5.43 | 5.75 |
| Dd2 IleRS V500A | Ile-tRNA synthetase | PF3D7_1343700 | 17.98 | 14.06 |
| Dd2 kelch13 C580C | Kelch protein K13 | PF3D7_1343700 | 15.16 | 13.94 |
| Dd2 kelch13 C580Y | Kelch protein K13 | PF3D7_0908800 | 6.36 | 4.72 |
| Dd2 kelch13 R539T | Kelch protein K13 | PF3D7_0523000 | 0.81 | 0.89 |
| Dd2 MDR1 F1072L | Multidrug resistance protein 1 | PF3D7_0509800 | 0.07 | 0.05 |
| Dd2 PI4K CNV | Phosphatidylinositol 4-kinase | PF3D7_0509800 | 2.48 | 2.59 |
| Dd2 PI4K S1320L+L1418F | Phosphatidylinositol 4-kinase | PF3D7_0509800 | 7.22 | 5.9 |
| Dd2 PI4K S743F+H1484Y | Phosphatidylinositol 4-kinase | PF3D7_1213800 | 1.92 | 2.37 |
| Dd2 ProRS L482H | Pro-tRNA synthetase | PF3D7_1011400 | 0.42 | 0.66 |
| Dd2 UDP-GT F37V | UDP-galactose transporter | PF3D7_1017000 | 0.05 | 0.03 |

**Supplementary Table S5.** Conditional knockdown (cKD) data.

| **cKD Line[a]** | | Reduced expression (–Shield-1) $IC_{50}$ (nM) | Full expression (+Shield-1) $IC_{50}$ (nM) |
|---|---|---|---|
| *Pf*PDK1 | Rep1 | 77.1 | 63.7 |
| | Rep2 | 58.8 | 58.1 |
| | Rep3 | 60.4 | 73.4 |
| | **Mean $IC_{50}$ ± SEM** | **65 ± 8** | **65 ± 6** |
| *Pf*NF45 control | Rep1 | 28.2 | 24.7 |
| | Rep2 | 45.0 | 19.2 |
| | Rep3 | 34.3 | 33.4 |
| | **Mean $IC_{50}$ ± SEM** | **36 ± 7** | **26 ± 6** |
| **cKD Line[b]** | | Reduced expression (1 nM aTc) $IC_{50}$ (nM) | Full expression (500 nM aTc) $IC_{50}$ (nM) |
| *Pf*Ark1 | Rep1 | 2.4 | 77.0 |
| | Rep2 | 2.8 | 54.3 |
| | Rep3 | 2.4 | 61.7 |
| | **Mean $IC_{50}$ ± SEM** | **2.5 ± 0.2** | **64 ± 9** |
| *Pf*Ark2 | Rep1 | 26.6 | 76.1 |
| | Rep2 | 22.1 | 65.0 |
| | Rep3 | 21.1 | 58.8 |
| | **Mean $IC_{50}$ ± SEM** | **23 ± 2** | **67 ± 7** |
| *Pf*PKAc | Rep1 | 32.0 | 99.7 |
| | Rep2 | 24.8 | 102.0 |
| | Rep3 | 21.4 | 85.4 |
| | **Mean $IC_{50}$ ± SEM** | **26 ± 4** | **95 ± 7** |
| *Pf*PI4K | Rep1 | 49.1 | 129.0 |
| | Rep2 | 59.2 | 92.0 |
| | **Mean $IC_{50}$ ± SEM** | **54 ± 7** | **110 ± 26** |
| **cKD Line[c]** | | Reduced expression (2.5 mM GlcN) $IC_{50}$ (nM) | Full expression (PBS vehicle) $IC_{50}$ (nM) |
| GSK3β-GFP-glmS | Rep1 | 70.9 | 74.5 |
| | Rep2 | 70.3 | 74.7 |
| | Rep3 | 70.4 | 74.6 |
| | **Mean $IC_{50}$ ± SEM** | **70.5 ± 0.2** | **74.58 ± 0.04** |
| **cKD Line[d]** | | Reduced expression (250 nM rapalog) $IC_{50}$ (nM) | Full expression (RPMI medium) $IC_{50}$ (nM) |
| GSK3β-DiCre | Rep1 | 85.5 | 75.4 |
| | Rep2 | 86.8 | 73.7 |
| | Rep3 | 86.7 | 74.2 |
| | **Mean $IC_{50}$ ± SEM** | **86.3 ± 0.4** | **74.5 ± 0.5** |

[a] Using the method by Hitz *et al.*[61]
[b] Using the method by Goldfless *et al.*[62]
[c] Using the method by Burda et al.[63]
[d] Using the method by Jones et al.[64]

**Supplementary Table S6**. List of oligonucleotides for donor vector construction.

| Description | Nucleotide sequence |
|---|---|
| Ark1 RHR forward sequence | aacccggaattcgagctcggGGTTTCACAACAAATGACATA |
| Ark1 RHR reverse sequence | cgagagattgggtattagacctagggataacagggtaatGAAGATGAGTTTAGTGAGGATA |
| Ark1 sgRNA target site | TCTTACGCGACACAACCTAG |
| Ark2 RHR forward sequence | aacccggaattcgagctcggACAATTCATATAAAAGAAAGCG |
| Ark2 RHR reverse sequence | cgagagattgggtattagacctagggataacagggtaatCATGTATCATTACTACATAAACAAGA |
| Ark2 sgRNA target site | GGAGTTAATATTAAAAGTAA |
| PKAc RHR forward sequence | aacccggaattcgagctcggTGATTGGTAGAAAAATGAAGAGA |
| PKAc RHR reverse sequence | agagattgggtattagacctagggataacagggtaatTTTTTATTAAGGTAATACTTAACATATA ACATTTTC |
| PKAc sgRNA target site | GGTTCATTCGCATAAAAAGG |

RHR, right homology region; cKD, conditional knockdown

**Ark1 LHR and re-codonized region (944-1549 bp; stop codon removed)**

CATTTGGATTTGAAACCCGAAAATGTTTTAGTAAACCATGAAGAAAAATGTAAGTTAGCTGATTTTGGTTTATC
AGCACATATAGGATCTAAACATAAAAAGAAAGGTATATCTCATTATAGAGGAACACATGATTATTGGTCTCCT
GAACAATGTGCAAGACATCAAAAGAAGAAACAGAATTTTGGAGAATTTGATCAGAAAACAGATATATGGACTC
TAGGGATTTTAGCTTTTGAATTAAAATTTGGTAGACCTCCATTTGGTTCAACAAATGAAGAAAGAGAAAATGTA
ATTATGAATAGAATACAAGATTATCATTGGAGTCAATTATTCTGTGAAAAAGTAAAACAAGATTTAATAGATAA
ATTATCACCTGAATTTAAAGATTTCTTAAATTTATGTCTAGATAAAAATCCAAAAAAAAGACCAACTGCAGAAT
CCTTAATTCAACATCCTTTCATCTTCATCCACAACAAGAACCGCTCCTGCATCAAGAGCTACGCCACCCAGCC
CCGCGGAAAGCAGGGCCCGAAGCTGAGCCACAAGGGCTTCAACGAGCAGGACGGCTCCTTCCTGACCCCC
ATCTGGTTCCACAATAAG

**Ark2 LHR and re-codonized region (4596-5468 bp; stop codon removed)**

ACTAGCCGACTTTGGATTTTCTTGTCAGCTCAAAAATAAAAGACAAAAAAGGAGTACATTCTGTGGCACAATC
GATTATATGCCCCCTGAAATTATTAACCAAATACCTTATGATTGTAATGTGGATCTGTGGTGTTTGGGTATAGT
TATATTTGAGTTGTTAGTTGGATTTCCTCCATTTACCGACGATACACAGgtaaaaaaataaaataaaaaaaaaaaaaattca
tatgtaattgaaatgaaataaaaacagtatgacatgtgcatataaaatggaataaaaaatatgacttgttcatgtaaaaaaaaaaaaaaaaaatatatatatata
tatatatatatatataaaaataacaaataaaataacatataaaattgctgacttgtacatgtaaaaaaaaaaaaaaaaaatatatatatatatatatatatataa
aaataacaaataaaataacatataaaattgctgacttgtacatgtgaaaactaccaattccaaaaagaacctttaaaaagacatttatgtaataataatatattattt
gatttattatatatgtattttttttttttttttagGAGAGGATCTTTGATCAGATCAAGGAGCTGAACTTCCACTTCCCCAAGAGCGT
GAGCCTGCTGGCCCAGGAGCTGATTCTGAAGCTGTGCAGCCGCACCGCCGAGGAGCGCATCAGCGCCGAC
GAGGTGAAGAGCCACCCGTGGATCAAGCAGTTCATC

**PKAc LHR and re-codonized region (975-1914 bp; stop codon removed)**

tttacacatattcattcataattccacttcaactttataccttttttatttttgaagAGATTTGAAACCTGAAAATTTATTACTTGATAAAGATGG
ATTTATAAAAATGACTGATTTTGGATTTGCTAAAATAGTCGAGACGAGAACTTATACTTTATGTGGAACTCCAG
AATATATCGCTCCAGAAATTTTATTGAACGTCGGACATGGAAAAGCCgtatgcaaaaaaaaaaaatgtggaaaaattacatga
acatatgcatatttatattgaaaagtgtgaatttattacatgaacatgttcataatttttatgaaaaaatgtggactattacatgaacatgttcataaattttatgaaaaaa
tgtggactattacatgaacatgttcataatttttatgaaaaaatgtggattattacatgatcattttatatatatatatatatatatatatatatatgtatatatttttatatttat
atgtttatatcaataggtttatcaaagaaatataaaatttttatttttacattatagGCTGATTGGTGGACTTTAGGTATATTTATCTATGAGAT
CCTTGTGGGTTGCCCACCATTCTACGCTAACGAGCCACTTCTTATATACCAAAAGATCCTTGAGGGGATCAT
CTACTTTCCAAAGTTCCTTGACAATAACTGTAAGCACCTTATGAAAAAGCTTCTTAGCCATGACCTTACCAAG
CGTTACGGGAACCTCAAAAAGGGTGCACAGAACGTGAAGGAGCATCCTTGGTTCTCAAACATAGACTGGGT
TAACCTTCTTAACAAGAACGTGGAAGTGCCTTACAAGCCAAAGTACAAGAACATATTCGACAGTAGTAACTTC
GAACGTGTTCAGGAGGACCTTACGATCGCAGACAAGATCACCAACGAGAACGACCCTTTCTACGACTGG

### 3. Methods

#### 3.1 Kinobeads screening

The assay including MS analysis was performed as described in Arendse *et al.*[60] Briefly chemoproteomic affinity capture experiments were performed as described previously[65] using Kinobeads,[66] which consist of promiscuous kinase inhibitors immobilized on Sepharose beads. Beads were equilibrated in lysis buffer and incubated at 4°C for 1 h with *P. falciparum* blood-stage protein extracts (*Pf*3D7 isolate; 0.3 mg), pretreated with either compound or DMSO (vehicle control). Protein depletion and $IC_{50}$ values were calculated using a TMT 10-plex setup,[67] enabling simultaneous relative quantification across 10 conditions. Apparent dissociation constants ($K_d^{app}$) were determined by accounting for protein depletion during rebinding experiments (**Supplementary Table S1**). Proteins were processed using a modified SP3 protocol,[68] labeled with TMT10 reagents, and subjected to LC-MS/MS analysis on Q Exactive Orbitrap or Orbitrap Fusion Lumos mass spectrometers (Thermo Fisher Scientific.[65, 69] Protein identification was performed using Mascot 2.4 (Matrix Science) with <1% false discovery rate (FDR) for single-spectrum assignments and <0.1% FDR for multi-spectrum assignments. Quantified proteins required ≥2 unique peptide matches with <0.1% FDR. Raw data tables are available in the **Supplementary Table S1** MS Excel file.

#### 3.2 *Plasmodium falciparum* parasitology methods

##### *3.2.1. Parasite culture*

Human malaria parasites were cultured as described previously with minor modifications.[70] Several culture-adapted strains of *P. falciparum* were used for the different assessments. Parasites were maintained at 5-10% parasitemia in human erythrocytes (type O+) suspended in RPMI-1640 (10.44 g/L) growth media supplemented with 25 mM HEPES, 4 g/L glucose, 0.088 g/L Hypoxanthine, 25 mM bicarbonate and 0.5% Albumax-II. Cultures were incubated at 37°C in a mixture of 3% $O_2$ and 4% $CO_2$ in nitrogen and had growth media replenished daily to ensure viability.

##### *3.2.2 Antiplasmodial activity assessments*

For the initial screen, the *in vitro* sensitivity of *Plasmodium falciparum* asexual blood stage parasites (3D7 strain) to the human kinase inhibitor set, including CHIR-124, was evaluated using a whole-cell [$^3$H]-hypoxanthine incorporation assay.[71] Compounds were tested in triplicate via a serial dilution series starting at 10 µM with a 1:3 dilution factor. Parasite growth inhibition was calculated relative to untreated controls, and $IC_{50}$ values were determined by fitting dose-response curves. Each assay plate included appropriate growth, vehicle, and positive control wells.

For follow up studies, CHIR-124 was commercially obtained from Selleckchem (Catalog No.S2683, SMILES: C1CN2CCC1C(C2)NC3=C(C(=O)NC4=C3C=C(C=C4)Cl)C5=NC6=CC=CC=C6N5, CAS No. 405168-58-3). A full dose-response assessment was performed for CHIR-124 in a 96-well plate to determine the concentration inhibiting 50% of parasite growth ($IC_{50}$ value), with parasite survival measured using lactate dehydrogenase (pLDH) activity.[72] Samples were prepared to a 10 mmol/L stock solution in 100% DMSO and stored at room temperature until testing. Dilutions to the desired starting concentration were freshly prepared in growth media on each occasion of the experiment The standard antimalarial drugs chloroquine and artesunate were used as the reference drugs in all experiments. The highest concentration of solvent to which the parasites were exposed was <0.5% and has no measurable effect on the parasite viability. The assay plate was incubated at 37 °C for 72 h in a sealed gas chamber under 3% $O_2$ and 4% $CO_2$ in nitrogen.

For activity assessed using the pLDH method, the wells in the assay plate were gently resuspended after 72 h, and 15 µL from each well was transferred to a corresponding well in a duplicate plate containing 100 µL of Malstat reagent and 25 µL of nitroblue tetrazolium solution. Plates were left to develop for 20 minutes in the dark and then absorbance of each well was quantified using a spectrophotometer at 620 nm wavelength. Regression analysis was performed using the Dotmatics software platform to quantify the $IC_{50}$ value.

#### 3.2.3. Stage specificity

Stage specificity assessments were carried out as described previously, using the pLDH assay as above with minor modifications.[73] Parasites at each stage were exposed to a range of drug concentrations from 6 μmol/L to 12 nmol/L and growth at each concentration compared between stages. Assessment of ring stage activity was carried out using a synchronous ring-stage culture at 2% parasitemia. Growth was determined via pLDH activity after 24h in the late trophozoite stage when pLDH activity is greatest. Schizont stage assessments were carried out using synchronous schizont cultures, and growth was determined after 48h in the next late trophozoite stage. Comparison of the relative parasite survival at each concentration between the two stages showed when the killing action was taking place.

#### 3.2.4 Stage-specific gametocyte production and luciferase assay

Gametocytogenesis was induced from a tightly synchronised (>97 % rings, 0.5 % parasitemia, 6 % hematocrit grown in RPMI media containing Albumax II in A+/O+ human erythrocytes) asexual parasite culture from a NF54-Pfs16-GFP-luc *P. falciparum* transgenic line, under stationary, hypoxic conditions as described before.[74] The hematocrit was reduced to 4 % after on day 0 as an anemic stressor and immature gametocytes (stage II/III) were harvested on day 5-6 from cultures exposed to 50 mM N-acetyl glucosamine (NAG) on days 1-4. For late-stage (stage IV/V) and/or mature (stage V) gametocytes, NAG treatment occurred from days 3-7, and gametocytes were harvested on day 10 or 13.

Compounds were evaluated for stage-specific gametocytocidal activity with a 48 h drug pressure on either immature (stage II/III) or mature (stage V) gametocytes (2 % gametocytemia, 1.5 % hematocrit) under hypoxic conditions at 37 ºC. Luciferase activity was determined as a proxy for viability as described before.[74] Effective half-maximal inhibition ($IC_{50}$) values were determined for 2-fold dilutions over 9 concentration points, in technical triplicates and for three independent biological replicates, with methylene blue and MMV390048 serving as internal controls. $IC_{50}$ values were determined through non-linear, 4-parameter curve fitting in GraphPad and are indicated as means ± S.E.

#### 3.2.5 Liver-stage assay

*Plasmodium berghei* (Pb) sporozoites were obtained by dissecting the salivary glands of infected *Anopheles stephensi* mosquitoes. These were provided by The SporoCore, at the University of Georgia, GA, USA (SporoCore.uga.edu). The parasites utilized a GFP-$Luc_{ama1-eef1}$ reporter line.[75] These engineered parasites, termed Pb-Luc, were used to infect HepG2-A16-CD81EGFP cells.

HepG2-A16-CD81EGFP cells stably transformed to express a GFP-CD81 fusion protein[76] were cultured at 37°C in 5% $CO_2$ in DMEM (Invitrogen, Carlsbad, USA) supplemented with 10% FBS (Corning, NY, USA), 1X Pen/Strep/Glu (Thermo Fisher Scientific, USA).

Compounds, including CHIR-124 (Selleckchem, Catalog No.S2683) and controls were prepared as 10 mM solutions in DMSO, and 10 nl of each (resulting in a final DMSO concentration of 0.1% per well) was transferred into assay plates using an ECHO 650 (Beckman). Concentrations ranged from 10 μM to 0.5 nM. Atovaquone (1 μM) served as a positive control, while 0.1% DMSO was used as a negative control. Human hepatic cells (HepG2-A16-CD81-EGFP) suspended in 5 μL of DMEM medium ($2x10^5$ cells/ml, supplemented with 5%FBS, 5XPen/Strep/Glu) were seeded (at $3 \times 10^3$ per well) in 1536-well plates (Greiner BioOne) 20 hours prior to infection. PbLuc sporozoites, isolated from the salivary glands of *A. stephensi* mosquitoes were filtered twice through a 20 μm nylon pore cell strainer. The sporozoites were resuspended in screening media, counted with a hemocytometer, and diluted to a final concentration of 200 sporozoites per μL. Each well received 1,000 sporozoites in 5 μL, and the plates were centrifuged for 3 minutes in an Eppendorf 5810 R (330 RCF) on the lowest acceleration and brake settings. After incubation at 37°C in 5% $CO_2$ for 48 hours, media was removed by centrifuging the inverted plates at 130 RCF for 1 minute. Wells then received 2 μL of Bright-Glo™ Luciferase Assay System (Promega).[77] Luminescence was measured immediately using a Pherastar FSX reader (BMG Labtech). Luminescence values were normalized using the positive and negative controls. $EC_{50}$ values were calculated with CDD Vault (Burlingame, CA). All experiments were performed with at least four technical replicates and repeated three times.

#### 3.2.6 *Liver cytotoxicity assay*

Plates were prepared following the liver stage conditions, except that 5 µl of the media was used instead of sporozoites.[77] After incubation at 37°C for 48 hours, media was removed by spinning the inverted plates at 130 RCF for 1 minute, after which 2 µl of CellTiter-Glo® Luminescent Cell Viability Assay (Promega) was dispensed in each well. Immediately after adding the CellTiter-Glo reagent, the plates were read by the Pherastar FSX reader (BMG Labtech). Luminescence intensity values were normalized against positive (Puromycin) and negative (DMSO) controls. $EC_{50}$ values were calculated in CDD Vault (Burlingame, CA). All experiments were performed in four technical replicates and repeated at least three times.

#### 3.2.7 P. falciparum *conditional knockdown assays*

The NF54 strain and PDK1 cKD parasites were cultured in the presence (control conditions) or absence of 675 nM Shield-1 (PfPDK1 knockdown conditions) as described by Hitz et al.[61] *In vitro* $IC_{50}$ determination of CHIR-124 against *P. falciparum* asexual blood stage (PDK1 cKD or control) parasites βwas performed with a modified [$^3$H]-hypoxanthine incorporation assay, as previously reported.[78]

Ark1 (PF3D7_0605300), Ark2 (PF3D7_0309200), and PKAc (PF3D7_0934800) cKD lines generated by fusing the coding sequence and non-coding RNA aptamer sequences in the 5'- and 3'-UTR, permitting translation regulation using the TetR-DOZI system.[79] Editing was achieved by CRISPR/SpCas9 using the linear pSN054 vector that contains cloning sites for the left homology region (LHR) and the right homology region (RHR) as well a gene-specific guide RNA under control of the T7 promoter. Constructs were clone into the pSN054 donor vector[80] that includes preinstalled V5-2xHA epitope tags, a 10x tandem array of TetR aptamers, and a multicistronic cassette for expression of TetR-DOZI (regulation), blasticidin S-deaminase (selection marker) and a Renilla luciferase (RLuc) reporter. The LHR and recoded region was installed in-frame with tandem V5-2x-hemagglutinin (HA) tags to afford C-terminal epitope-tagged proteins post-editing, and upstream of the regulatory aptamer array. All primer and synthetic fragment sequences generated using the BioXP™ system are included in **Supplementary Table S6**. The final construct was sequence-verified and further confirmed by restriction digests. Transfection into Cas9- and T7 RNA polymerase-expressing NF54 parasites was carried out by pre-loading erythrocytes with the donor vector as previously described.[81] Parasite culture was maintained continuously in 500 nM anhydrotetracycline (aTc, Sigma-Aldrich 37919) and drug selection with 2.5 µg/mL of Blasticidin S (RPI Corp B12150-0.1) was initiated four days after transfection. Cultures were monitored by Giemsa smears and RLuc measurements.

Compound susceptibility assays using *P. falciparum* Ark1, Ark2, PKAc and PI4K cKD lines were carried out as previously described.[79, 80] Briefly, synchronous ring-stage Ark1 (PF3D7_0605300), Ark2 (PF3D7_0309200), PKAc (PF3D7_0934800) and PI4K (PF3D7_0509800) cKD parasites, as well as a control parasite line expressing an aptamer-regulatable fluorescent protein were maintained in the presence of high aTc (500 nM) or no aTc and distributed into 384-well polystyrene microplates (Corning). Stock solutions of compounds were serially diluted and transferred to the parasite-containing plates using the Janus platform (PerkinElmer). DMSO and dihydroartemisinin treatment (500 nM) served as reference controls. Luminescence was measured after 72 h using the Renilla-Glo Luciferase Assay System (Promega E2750) and the GloMax Discover Multimode Microplate Reader (Promega), and $IC_{50}$ values were obtained from corrected dose-response curves using Graph-Pad Prism.

For the generation of the GSK3β cKD lines, parasite culture medium excluded glucose, B+ human erythrocytes were used and cultures were incubated with 1% $O_2$ and 5% $CO_2$. The lines were generated by selection linked integration (SLI) as described previously.[82] GSK3β-GFP-glmS[83] was generated by endogenous C-terminal GFP-tagging of the target gene thereby introducing a glmS ribozyme sequence[84] upstream of the 3' untranslated region as described previously.[63] GSK3β-DiCre was generated using a N-terminal homology region of the target gene followed by a recodonized version of the C-terminal kinase domain flanked by artificial introns containing loxP sites as described previously.[64] Subsequently, parasites were transfected with an episomal copy of a dimerizable Cre recombinase (DiCre) and excision of the GSK3β kinase domain was induced by rapalog-mediated dimerization of DiCre as described previously.[64]

Drug sensitivity assays with GSK3β-cKD lines were carried out as described previously).[85] Briefly, synchronous ring stage parasites were seeded in to black flat-bottom 96 well plates (ThermoFisher) at 0.1 % parasitemia and 2% hematocrit at a final volume of 200 µl. GlmS-based knock down and DiCre-based knockout were induced by treatment with 2.5 mM Glucosamine or 250 nM rapalog (Takara Clontech) respectively. Vehicle treated parasites served as a control. Compound at varying concentrations was added in triplicates and parasites were incubated for 96 h under standard culture conditions. Parasite proliferation was quantified based on the DNA content by replacing 100 µl of the supernatant with 1x SYBR Gold stain (Invitrogen) in lysis buffer (20 mM Tris-HCl pH 7.5; 5 mM EDTA, 0.008% saponin; 0.08% Triton-X-100). Fluorescence of SYBR-Gold-labelled DNA was quantified using an EnVision multilabel plate reader (PerkinElmer, 485 nm excitation, 535 nm emission) as described previously.[86] $IC_{50}$ values were calculated from dose-response curves using GraphPad Prism.

*3.2.8 Cross-resistance profiling using the Antimalarial Resistome Barcode sequencing assay (AReBar)*

Profiling of cross-resistance using a pooled library of barcoded mutant parasite lines was performed as previously described.[87, 88] The pool consisted of parasites in the *Pf*3D7 and *Pf*Dd2 backgrounds, with mutations described in **Supplementary Table S4**. The pool was exposed to either $3\times IC_{50}$ of CHIR-124, or untreated (control), and cultured for 14 days with growth measured by flow cytometry every 2-3 days. Parasitemia was maintained below 5%. PCR-amplified barcodes were quantified by next generation sequencing using an Oxford Nanopore minION. The log2 fold change (LFC) in proportion was calculated relative to the untreated control pool, with a LFC >2.5 indicating cross-resistance.

*3.2.9 Combination studies via fixed-ratio isobolograms*

Isobolograms were determined using the fixed-ratio isobole analysis method previously described.[89] This was carried out by treating intraerythrocytic ABS parasites with fixed drug combination ratios of CHIR-124 and chloroquine or hesperadin set at 5:0, 4:1, 3:2, 2:3, 1:4, and 0:5. Independent dose-response curves of the fixed drug combinations were determined using the pLDH assay against NF54 ABS parasites for 72 h. Fractional $IC_{50}$ values were calculated individually for each drug by plotting the relevant fractional concentration vs % parasite survival using GraphPad Prism v10. Mean fractional $IC_{50}$ values for each drug pair were plotted and summed to determine effects of synergy, additivity or antagonism.

*3.2.10 Rate- and stage-specific morphological evaluations and inhibitor effect reversibility*

To determine stage specificity during asexual intra-erythrocytic development, *P. falciparum* NF54 parasites were synchronised using 5 % D-Sorbitol to obtain a >90 % ring-stage population (6 – 10 h post-invasion, hpi). *In vitro*, synchronised parasite cultures were treated with CHIR-124 ($3\times IC_{50}$). The effect of CHIR-124 on various asexual intra-erythrocytic stage parasites (ring, early/late trophozoite, and schizont) was monitored morphologically every 12 h using Giemsa-stained thin smears. Images were captured using a Nikon Eclipse 50i light microscope adapted with a Nikon camera (Nikon DS-Fi1) and NIS-Elements software.

To further assess and quantify the specific parasite stage at which CHIR-124 is active and to determine whether the effect is reversible, as described above, various asexual intra-erythrocytic stages were treated with CHIR-124 at $3\times IC_{50}$ for 12-h segments. After which, CHIR-124 was washed off, and parasites were left to recover in drug-free medium. Parasitaemia and nuclear content were monitored every 12 h until the next ABS life cycle using flow cytometry, as described in *van Biljon R et al.*[90]

#### 3.3. β-hematin/hemozoin formation inhibition assays and metal chelation studies

*3.3.1 NP-40 based extracellular β-hematin formation inhibition assay*

The NP-40 detergent based assay methods for inhibitors of βH formation described by Sandlin *et al.*[91] were modified for manual liquid delivery. Samples were dissolved in DMSO to give 10 mM solutions and 20 µL of each was delivered to wells in the last column (column 12) of a 96-well plate together with distilled water (140 µL) and NP40 detergent (305.5 mM, 40 µL). A solution containing water/NP40 (305.5 mM)/DMSO at a v/v ratio of 70%/20%/10% respectively was prepared and then 100 µL was added to all other wells (columns 1–11). A serial dilution of each compound (100 µL) from column 12 down to column 2 was carried out. Column 1 served as a blank with 0 mM sample. A 25 mM hematin stock solution was prepared by sonicating hemin in DMSO for one minute and then suspending 178 µL of this in a 1 M acetate buffer (20 mL, pH 4.8). The homogenous suspension (100 µL) was then added to the wells to give final buffer and haematin concentrations of 0.5 M and 100 µM respectively. The plate was covered and incubated at 37 °C for 5–6 h in an incubator. Analysis was carried out using the pyridine-ferrichrome method developed by Ncokazi and Egan.[92] A solution of 50% (v/v) pyridine, 30% (v/v) $H_2O$, 20% (v/v) acetone and 0.2 M HEPES buffer (pH 7.4) was prepared and 32 µL added to each well to give a final pyridine concentration of 5% (v/v). Acetone (60 µL) was then added to assist with hematin dispersion. The UV-vis absorbance of the plate wells was read on a Thermo Scientific MultiskanGO plate reader. Sigmoidal dose-response curves were fitted to the absorbance data using GraphPad Prism v5 to obtain a 50% inhibitory concentration ($IC_{50}$) for each compound.

#### 3.3.2 *Cellular heme fractionation assay*

Heme fractionation analysis was performed as described.[93] Briefly, early ring-stage sorbitol- synchronized NF54 parasites were exposed to CHIR-124 at concentrations corresponding to 0.5, 1, 1.5, 2 and 3 times the ABS $IC_{50}$. After a 32 h incubation, cells were saponin-lysed and trophozoites were subjected to a series of lysis, solubilization, and differential centrifugation steps to obtain fractions that contain Hb, free heme or Hz. The UV-visible spectra of heme as Fe(III)heme-pyridine was analyzed in each of these fractions and compared to a heme standard curve for accurate quantification. These quantities were subsequently normalized using the number of analyzed cells, as determined by flow cytometry of an aliquot of exposed cells. Two-tailed t-tests were used to assess significance.

#### 3.3.3 *Metal chelation studies*

To construct Job plots to determine, qualitatively, the extent of chelation between CHIR-124 and physiologically-relevant metal ions, the absorption spectra of solutions of varying mole fractions of CHIR-124 and metal ions were recorded according to a method previously described, with modifications.[94] Briefly, the total concentration of the system was constrained to 0.3 mM and the absorbance was measured from 200 nm to 600 nm in a 1.00 mm path length cuvette in 20 mM PBS at pH 7.4. Working solutions with mole fractions of metal ion between 0 and 1 were prepared, i.e. at metal ion mole fractions of 0, 0.2, 0.4, 0.6, 0.8 and 1, with the remainder of the mole fraction provided by a complementary amount of CHIR-124 (1, 0.8, 0.6, 0.4, 0.2 and 0, respectively). Spectrophotometric data were analysed at the absorbance maximum of CHIR-124 (364 nm) and normalised according the maximum and minimum absorbance values recorded for each experiment.

### 3.4 Biochemical assays

#### 3.4.1 *KinaseSeeker assay*

KinaseSeeker is a homogeneous competition binding assay where the displacement of an active site-dependent probe by an inhibitor is measured by a change in luminescence signal. Luminescence readout translates into a highly sensitive and robust assay with low background and minimal interference from test compounds.[95, 96]

10 mM stock solutions of compounds were serially diluted to obtain assay stocks. Prior to initiating a profiling campaign, the compounds were evaluated for false positive against split-luciferase. The compounds were then screened in duplicate against each of the kinases. For kinase assays, each Cfluc-Kinase was translated along with Fos-Nfluc using a cell-free system (rabbit reticulocyte lysate) at 30 ˚C for 90 min. 24 µL aliquot of this lysate containing either 1 µL of DMSO (for no-inhibitor control) or compound solution in DMSO (various concentrations) was incubated for 2 hours at room temperature in presence of a kinase specific probe. 80 µL of luciferin assay reagent was added to each solution and luminescence was measured on a luminometer.

The % activity remaining was calculated using the following equation:

$$\% \text{ Inhibition} = \frac{\text{ALU}_{\text{Control}} - \text{ALU}_{\text{Sample}}}{\text{ALU}_{\text{Control}}} \times 100$$

$$\% \text{ Activity Remaining} = 100 - \% \text{ Inhibition}$$
